# A neuropeptide-specific signaling pathway for state-dependent regulation of the mesolimbic dopamine system

**DOI:** 10.1101/2025.07.17.665396

**Authors:** Mollie X. Bernstein, Omar Koita, Marta Trzeciak, Andrew Fan, Daniel T. McAuley, Seung-Woo Jin, Larry S. Zweifel

## Abstract

Dopamine (DA)-producing neurons of the ventral tegmental area (VTA) regulate consummatory behavior in a state-dependent manner (e.g. when hungry or thirsty). The mechanisms by which and extent to which DA neurons are regulated by these interoceptive signals are poorly understood. Here, we identify transient receptor potential canonical type 6 (TRPC6) channels as selective mediators of neuropeptide receptor-induced calcium signaling in VTA-DA neurons. These channels regulate DA neuron activity and consummatory behavior in a state-dependent manner. We find that TRPC6 channels regulate distinct aspects of neuropeptide-induced calcium signals in DA neurons but make little contribution to calcium dynamics associated with metabotropic neurotransmitter receptor signaling. We further show that TRPC6 channels regulate scalable reward valuation and consummatory behavior in hungry but not thirsty mice. These findings demonstrate that neuropeptide-and neurotransmitter-activated G-protein coupled receptors (GPCRs) regulate cellular calcium dynamics through distinct mechanisms, and that TRPC6 channels are important determinants of how animals respond to different homeostatic demands.

## INTRODUCTION

Midbrain dopamine (DA)-producing neurons of the ventral tegmental area (VTA) play a critical role in scaling responses to rewards and regulating reward-seeking behavior (Wise, 2004), as well as in homeostatically driven fluid and food consumption (Mietlicki-Baase et al., 2021; Palmiter, 2007). Numerous neuropeptides that regulate energy balance and homeostatic fluid and food consumption modulate the activity of VTA DA neurons (Khan et al., 2024; Naef et al., 2015; Palmiter, 2007), and several of these have been shown to increase DA neuron activity and dopamine release (Cone et al., 2014; Nalivaiko et al., 1997; Seutin et al., 1989). These stimulatory neuropeptides typically signal through GPCRs that are coupled to G_q/11_-mediated signaling cascades to increase intracellular calcium, but the mechanisms through which they modulate VTA-DA neurons are largely unknown.

The two most abundantly enriched stimulatory neuropeptide receptors in VTA-DA neurons are the neurotensin receptor (Ntsr1) and the neurokinin B (also known as tachykinin 2) receptor (Tacr3) (Chung et al., 2017). Ntsr1 and Tacr3 are stimulatory Gα_q/11_-protein coupled receptors (GPCRs) and upon ligand binding, phospholipase C (PLC) is activated, which cleaves PIP_2_ into IP_3_ and diacylglycerol (DAG). Generation of IP_3_ activates IP_3_ receptors on the endoplasmic reticulum to induce intracellular calcium release (Clapham, 1995). DAG, in turn, activates protein kinase C (PKC), which then phosphorylates other proteins, leading to multiple cellular responses, including the phosphorylation of ion channels to regulate neuronal excitability (Kaczmarek, 1987).

PIP_2_ is a critical co-factor for the Kv7 family of voltage-gated potassium channels; thus, PIP_2_ hydrolysis also results in membrane depolarization (Suh and Hille, 2002) which can, in turn, activate voltage-gated calcium channels (VGCCs) (Mathes and Thompson, 1994).

An additional mechanism for the regulation of extracellular calcium influx and the modulation of neuronal excitability through neuropeptide signaling is the transient receptor potential canonical (TRPC) channels (Kelly et al., 2018; Qiu et al., 2021; Stuhrman and Roseberry, 2015). TRPC channels are a class of calcium-permeable ion channels (Wang et al., 2020), subdivided into 3 subtypes (TRPC1/4/5, TRPC2, and TRPC3/6/7) based on their sequence homology and functional similarities (Wang et al., 2020; Zhang et al., 2023). In *Kisspeptin* (*Kiss1*)-expressing neurons in the arcuate nucleus (ARH), activation of the Tac2/NkB receptor Tacr3 results in a TRPC5-dependent membrane depolarization (Qiu et al., 2016). Tac2/NkB along with the co-release of dynorphin, coordinates the synchronous firing of ARH-*Kiss1* neurons (Kelly and Wagner, 2024).

Given the high expression of *Tacr3* in VTA-DA neurons, we hypothesized that Tacr3-dependent activation of a TRPC channel is an important regulatory pathway for neuropeptide modulation of VTA-DA neurons. Of the TRPC channels, the receptor operated channel TRPC6 is highly enriched in DA neurons (Chung et al., 2017) and is the most abundantly expressed TRPC channel in these cells (Wang et al., 2024). Using a cell-type-specific CRISPR/Cas9 mutagenesis approach, we demonstrate that TRPC6 channels are selectively involved in rapid neuropeptide receptor-induced calcium influx, but not neurotransmitter receptor-induced calcium increases. We further find that TRPC6 channels regulate neuropeptide-induced synchronous calcium increases and calcium oscillations in VTA dopamine neurons. During liquid sucrose consumption in thirsty or hungry mice, we find that genetic inactivation of *Trpc6* impairs scalar consummatory responses to increasing sucrose concentrations and scalar increases in calcium signals in VTA-DA neurons in hungry but not thirsty mice. These findings demonstrate that TRPC6 channels act downstream of stimulatory neuropeptide receptor signaling and modulate consummatory responses in a state-dependent manner.

## RESULTS

### CRISPR/SaCas9 mutagenesis of *Trpc6* in VTA-DA neurons

To confirm that *Trpc6* is the predominant TRPC channel expressed in DA neurons, we analyzed our previously published dataset of actively translated mRNA from VTA-DA neurons (Chung et al., 2017). Among the seven TRPC channel genes, *Trpc6* is the most abundantly expressed and is highly enriched in VTA-DA neurons relative to other VTA cell types (Fig. 1A), consistent with previous reports (Wang et al., 2024). RNAscope *in situ* hybridization analysis further demonstrated that *Trpc6* expression is localized to a large proportion of *Th*-positive cells (a marker for DA cells) and is relatively restricted to expression in VTA-DA neurons (Fig. 1B).

**Figure 1.**
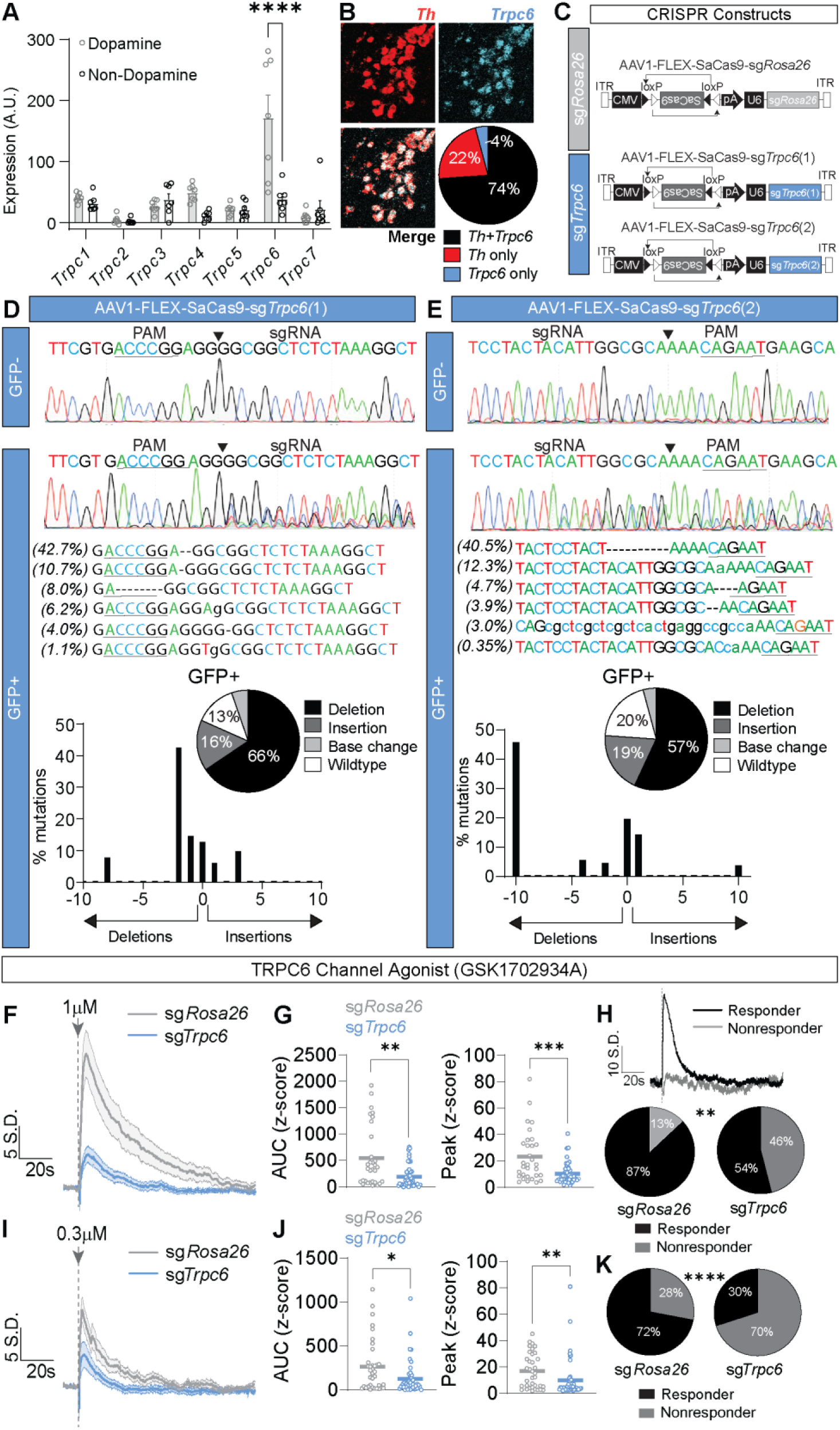
Validation of *Trpc6* mutagenesis approach. (A) mRNA expression of TRPC channel-encoding genes in the VTA of adult mice adapted from Chung et al. (2017) (N=7 mice; Two-way repeated measures ANOVA followed by Bonferroni post hoc comparison, ****P<0.0001). (B) *In situ* hybridization experiment showing localization of *Th, Trpc6,* and co-localization of *Th* and *Trpc6* (N=3 mice, n=3 hemi sections/mouse). (C) Schematic showing the CRISPR constructs injected into the sg*Rosa26* (control) and sg*Trpc6* (experimental) groups. (D, E) Quantification of mutagenesis in the *Trpc6* gene in VTA-DA GFP-and GFP+ cells in AAV1-FLEX-SaCas9-sg*Trpc6*(1) injected mice (N=3 mice) (D) and AAV1-FLEX-SaCas9-sg*Trpc6*(2) injected mice (N=3 mice) (E). Dashed lines indicate nucleotide deletions; lowercase letters indicate nucleotide insertions. (F-K) *Ex vivo* slice calcium imaging experiment of VTA-DA cells with a 50 ms puff application of TRPC6 channel direct agonist GSK1702934A at 1 µM (F-H) and 0.3 µM (I-K). Mean ± S.E.M. z-score traces (F, I), AUC and peak analyses for 1 µM (G) and 0.3 µM (J). (H, K) Responder classification analysis for both concentrations; example trace of both types are shown in the top panel of (H). (G-H) (N= 4 sg*Rosa26* mice, n=31 cells and N= 3 sg*Trpc6* mice, n=46 cells; D’Agostino & Pearson Test, Mann Whitney, Chi-squared test, **P<0.01, ***P<0.001). (J-K) (N= 3 sg*Rosa26* mice, n=36 cells and N= 3 sg*Trpc6* mice, n=47 cells; D’Agostino & Pearson Test, Mann Whitney, Chi-squared test, *P<0.05, **P<0.01, ****P<0.0001). See Extended Data Table 1 for details on statistical tests.

To investigate the function of TRPC6 channels in VTA-DA neurons, we selectively mutated the *Trpc6* gene using viral-mediated cell-type-specific CRISPR/Cas9 mutagenesis (Hunker et al., 2020). We designed two separate sgRNAs targeting different exons in the *Trpc6* gene, to generate two viral constructs: AAV1-CMV-FLEX-SaCas9-U6-sg*Trpc6*(1) and AAV1-CMV-FLEX-SaCas9-U6-sg*Trpc6*(2) (Fig. 1C). Deep sequencing of PCR amplicons containing the sgRNA targeted regions revealed a high degree of insertion and deletion mutations (indels) (Fig. 1D-E).

To confirm loss of TRPC6 channel function, we co-injected DAT-Cre mice with either AAV1-FLEX-sg*Rosa26* or a mixture of AAV1-FLEX-sg*Trpc6*(1) and AAV1-FLEX-sg*Trpc6*(2), along with the genetically encoded calcium indicator GCaMP8f into the VTA. We assessed calcium influx in VTA-DA neurons in acute brain slices following a brief (50 ms) focal application (puff) (Forman et al., 2017) of the TRPC6 channel agonist GSK1702934A.

A 50 ms puff resulted in large increases in calcium fluorescence in sg*Rosa26* control cells at a high (1 µM GSK1702934A) (Fig. 1F-G) and intermediate (0.3 µM GSK1702934A) concentration (Fig. 1I-J). These responses were significantly attenuated in sg*Trpc6* cells for both concentrations of GSK1702934A (Fig. 1F-G, I-J). VTA-DA neurons in the sg*Trpc6* group also displayed a significant reduction in the proportion of responsive and nonresponsive cells for both 1 µM GSK1702934A (Fig. 1H) and 0.3 µM GSK1702934A (Fig. 1K). Together, these data confirm the high functional efficiency of *Trpc6* mutagenesis in VTA-DA neurons.

### TRPC6 channels are selectively coupled to neuropeptide receptor activation

Based on the evidence linking TRPC5 channels to Tacr3 signaling (Kelly and Wagner, 2024) and the high expression of *Tacr3* in VTA-DA neurons (Chen et al., 1998; Chung et al., 2017), we first performed *ex vivo* calcium imaging of VTA-DA neurons using the Tacr3 agonist senktide. As above, DAT-Cre mice were bilaterally injected with either AAV1-FLEX-sg*Rosa26* or a combination of AAV1-FLEX-sg*Trpc6*(1) and AAV1-FLEX-sg*Trpc6*(2) along with AAV1-FLEX-GCaMP8f into the VTA. We observed a large increase in calcium fluorescence in response to a puff of 0.5 µM senktide, which was significantly reduced in the sg*Trpc6* group quantified as both AUC and peak amplitude (Fig. 2A). There was no significant difference in the proportion of responsive versus nonresponsive cells (Fig. 2A).

**Figure 2.**
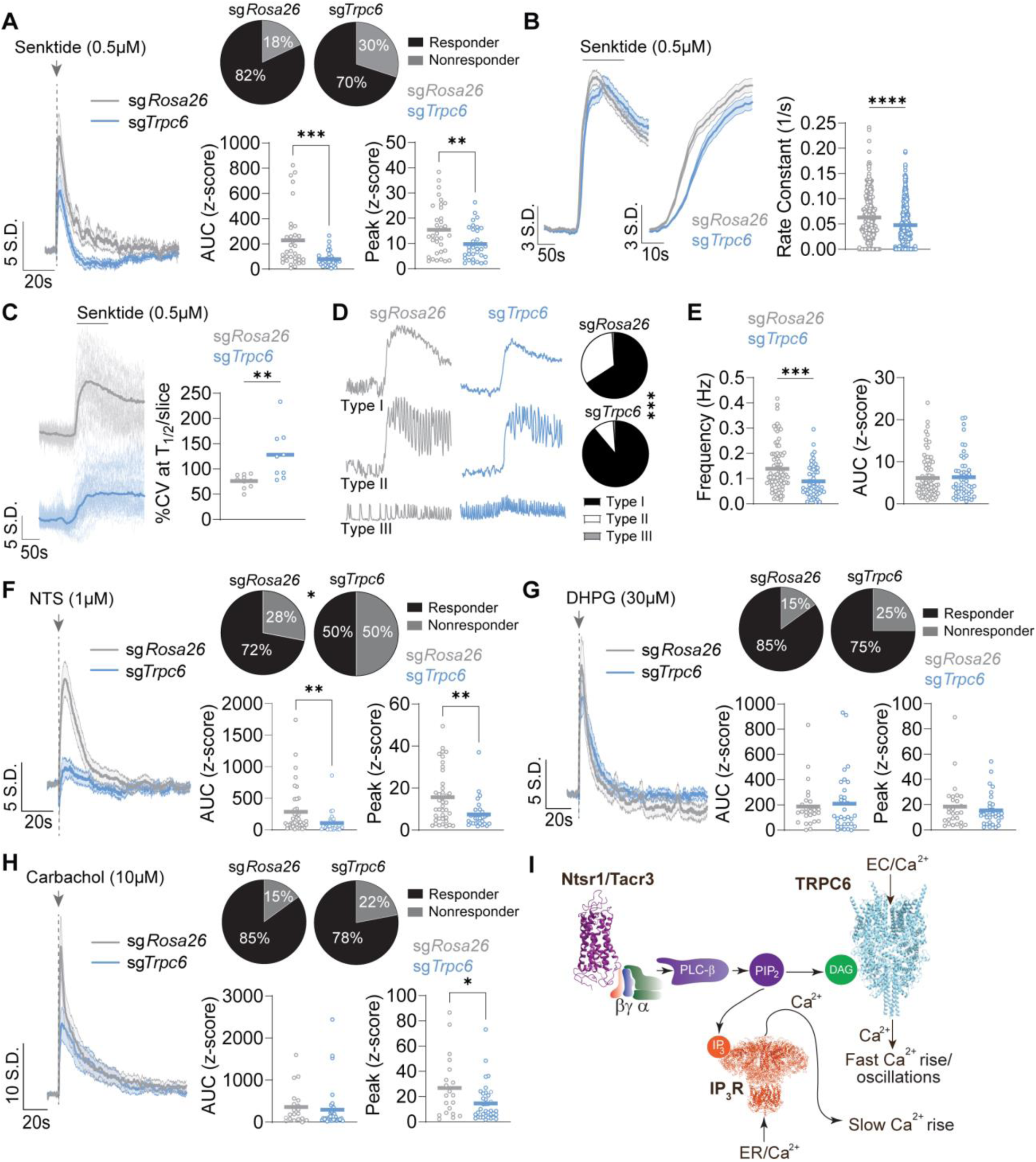
TRPC6 channels are coupled to neuropeptide receptor signaling pathways. (A) *Ex vivo* slice calcium imaging experiment of VTA-DA cells following a 50 ms puff application of senktide (0.5 µM) with mean ± S.E.M. z-score traces, AUC, peak, and responder classification analyses (N=4 sg*Rosa26* mice, n=34 cells and N=4 sg*Trpc6* mice, n=33 cells; D’Agostino & Pearson Test, Mann Whitney, Chi-squared test, **P<0.01, ***P<0.0001). (B-E) *Ex vivo* slice calcium imaging experiment of VTA-DA cells following a 5 min bath application of senktide (0.5 µM). (B) Average rise (left panel), initial rise (middle panel) and rate constant analysis (right panel) for the initial rise time of calcium fluorescence (N=3 sg*Rosa26* mice, n=290 cells and N=3 sg*Trpc6* mice, n=512 cells; Independent t-test, ****P<0.0001). (C) Data from one slice per group from (B) where the solid line is the average trace (left panel); coefficient of variation (CV), expressed as a percentage, taken at T_1/2_ per slice (right panel) (N=3 sg*Rosa26* mice, n=10 slices and N=3 sg*Trpc6* mice, n=9 slices; Independent t-test, **P<0.01). (D) Example traces for Type I, II, and III responsive cells (left panel) and proportion analysis (right panel) (Fisher’s exact test, ***P<0.001). (E) Frequency of oscillations for Type II and III cells (left panel) and AUC analysis (right panel) (N=3 sg*Rosa26* mice, n=84 cells and N=3 sg*Trpc6* mice, n=52 cells; D’Agostino & Pearson Test, Mann Whitney, ***P<0.001). (F) *Ex vivo* slice calcium imaging experiment of VTA-DA cells with a 50 ms puff application of neurotensin (NTS) (1 µM) with mean ± S.E.M. z-score traces, AUC, peak, and responder classification analyses (N= 3 sg*Rosa26* mice, n=40 cells and N= 3 sg*Trpc6* mice, n=30 cells; D’Agostino & Pearson Test, Mann Whitney, Chi-squared test, *P<0.05, **P<0.01). (G) *Ex vivo* slice calcium imaging experiment of VTA-DA cells with a 50 ms puff application of (S)-3,5-Dihydroxyphenylglycine (DHPG) (30 µM) with mean ± S.E.M. z-score traces, AUC, peak, and responder classification analyses (N= 3 sg*Rosa26* mice, n=27 cells and N=3 sg*Trpc6* mice, n=32 cells; D’Agostino & Pearson Test, Mann Whitney, Chi-squared test). (H) *Ex vivo* slice calcium imaging experiment of VTA-DA cells with a 50 ms puff application of carbachol (10 µM) with mean ± S.E.M. z-score traces, AUC, peak, and responder classification analyses (N=3 sg*Rosa26* mice, n=20 cells and N=4 sg*Trpc6* mice, n=37 cells; D’Agostino & Pearson Test, Mann Whitney, Chi-squared test, *P<0.05). (I) A model by which activation of neurotensin receptor 1 (Ntsr1) and tachykinin receptor 3 (Tacr3) initiates a signaling cascade resulting in extracellular (EC) Ca^2+^ ions flowing into the cell through TRPC6 channels contributing to a fast rise in [Ca^2+^] and Ca^2+^ oscillations. Ca^2+^ ions exit the endoplasmic reticulum (ER) via IP_3_ receptors giving rise to a slow rise in [Ca^2+^]. IP_3_ = inositol trisphosphate, PLC-β = phospholipase C beta, PIP_2_ = phosphatidylinositol bisphosphate, DAG = diacylglycerol. See Extended Data Table 1 for details on statistical tests.

In ARH-*Kiss1* neurons, bath application of senktide results in a TRPC5-dependent, STIM/ORAI-independent, augmentation of the synchronous calcium rise in these cells (Qiu et al., 2021). To assess whether TRPC6 mediates a similar function in VTA-DA neurons, we bath applied senktide (500 nM) to acute brain slices from DAT-Cre mice injected with either AAV1-FLEX-sg*Rosa26* or AAV1-FLEX-sg*Trpc6*(1) and AAV1-FLEX-sg*Trpc6*(2) along with AAV1-FLEX-GCaMP8f into the VTA. Both groups exhibited large increases in calcium fluorescence following senktide application (Fig. 2B). Consistent with TRPC6-dependent rapid calcium increases associated with brief, focal senktide application (Fig. 1A), the initial rate of rise (1/s) was significantly slower in sg*Trpc6* cells compared to control sg*Rosa26* cells (Fig. 2B). To determine whether TRPC6 channels facilitate the synchronous rise in calcium following Tacr3 activation, we assessed the variance (percent coefficient of variation around the mean rise at T_1/2_) of calcium increases within a slice. The variance in sg*Trpc6* cells was significantly increased compared to sg*Rosa26* cells (Fig. 2C).

During our analysis, we observed three distinct responses in calcium signals in VTA-DA neurons to senktide application. The largest number of cells (Type I) displayed a sustained increase in calcium, followed by cells with a sustained increase coupled with oscillatory calcium signals (Type II), and a rarer group of cells (∼1%) which showed intrinsic calcium oscillations enhanced by senktide (Type III) (Fig. 2D). Mutagenesis of *Trpc6* resulted in a significant reduction in the number of oscillating cells induced by senktide compared to controls (Fig. 2D). Of the cells in which senktide-induced oscillations were observed, the frequency of oscillations was significantly reduced in sg*Trpc6* cells compared to sg*Rosa26* cells (Fig. 2E); however, the amplitude of the oscillation was unaffected (Fig. 2E).

In addition to *Tacr3,* the NTS receptor NTSR1 (*Ntsr1*) is highly expressed in VTA-DA neurons and facilitates a robust increase in intracellular calcium *in vivo* (Soden et al., 2023). Application of a non-selective TRPC channel blocker also attenuates the neurotensin-mediated inward current in VTA-DA neurons (Stuhrman and Roseberry, 2015). To assess whether TRPC6 acts downstream of NTSR1 activation, we focally applied NTS (1 µM) as above (50 ms puff). NTS-evoked calcium was significantly reduced in sg*Trpc6* cells as quantified by AUC and peak amplitude compared to sg*Rosa26* cells (Fig. 2F).

In other cell types, TRPC6 channels are downstream of other types of GPCRs that cause production of DAG, such as metabotropic glutamate receptors (mGluR) (Wang et al., 2019) and muscarinic acetylcholine receptors (mAChR) (Zhang et al., 2006). To determine whether TRPC6 channels are general downstream effectors of G_q/11_-coupled receptors, we focally applied the mGluR agonist (S)-3,5-Dihydroxyphenylglycine (DHPG) and the mAChR agonist carbachol to VTA-DA neurons. Both DHPG (30 µM) and carbachol (10 µM) induced rapid and robust increases in intracellular calcium (Fig. 2G-H). *Trpc6* mutagenesis did not affect DHPG-evoked calcium signals in VTA-DA neurons (Fig. 2G) and had only a modest but significant effect on the peak carbachol-evoked calcium signal, without affecting the overall calcium signal (Fig. 2H). These results suggest that TRPC6 channels act predominantly downstream of neuropeptide receptor activation (Fig. 2I).

### *Trpc6* mutagenesis does not alter spontaneous or evoked action potential firing in VTA-DA neurons

Previous findings have shown that *Trpc6* knockdown in VTA-DA neurons leads to reduced neuronal intrinsic excitability and resting membrane potential (Wang et al., 2024). However, others have shown that mutagenesis of *TRPC6* in human induced pluripotent stem cell-derived pyramidal neurons causes an increase in neuronal excitability (Shin et al., 2023). To determine whether CRISPR/SaCas9 mutagenesis of *Trpc6* in adult VTA-DA neurons alters neuronal excitability, we performed whole-cell patch-clamp recordings in acute brain slices from DAT-Cre mice injected with either AAV1-FLEX-sg*Rosa26* or AAV1-FLEX-sg*Trpc6*(1) and AAV1-FLEX-sg*Trpc6*(2) along with AAV1-FLEX-GcaMP8f (Fig. 3A). We did not observe any significant differences in the proportion of spontaneous versus inactive cells, firing rate, or CV-ISI of spontaneously active cells (Fig. 3B-D). Additionally, we did not observe any differences in resting membrane potential or evoked excitability, as measured by a ramp protocol and current injection (Fig. 3E-H). These results indicate that *Trpc6* loss of function in VTA-DA neurons does not result in an overt change in the intrinsic electrophysiological properties of these cells.

**Figure 3.**
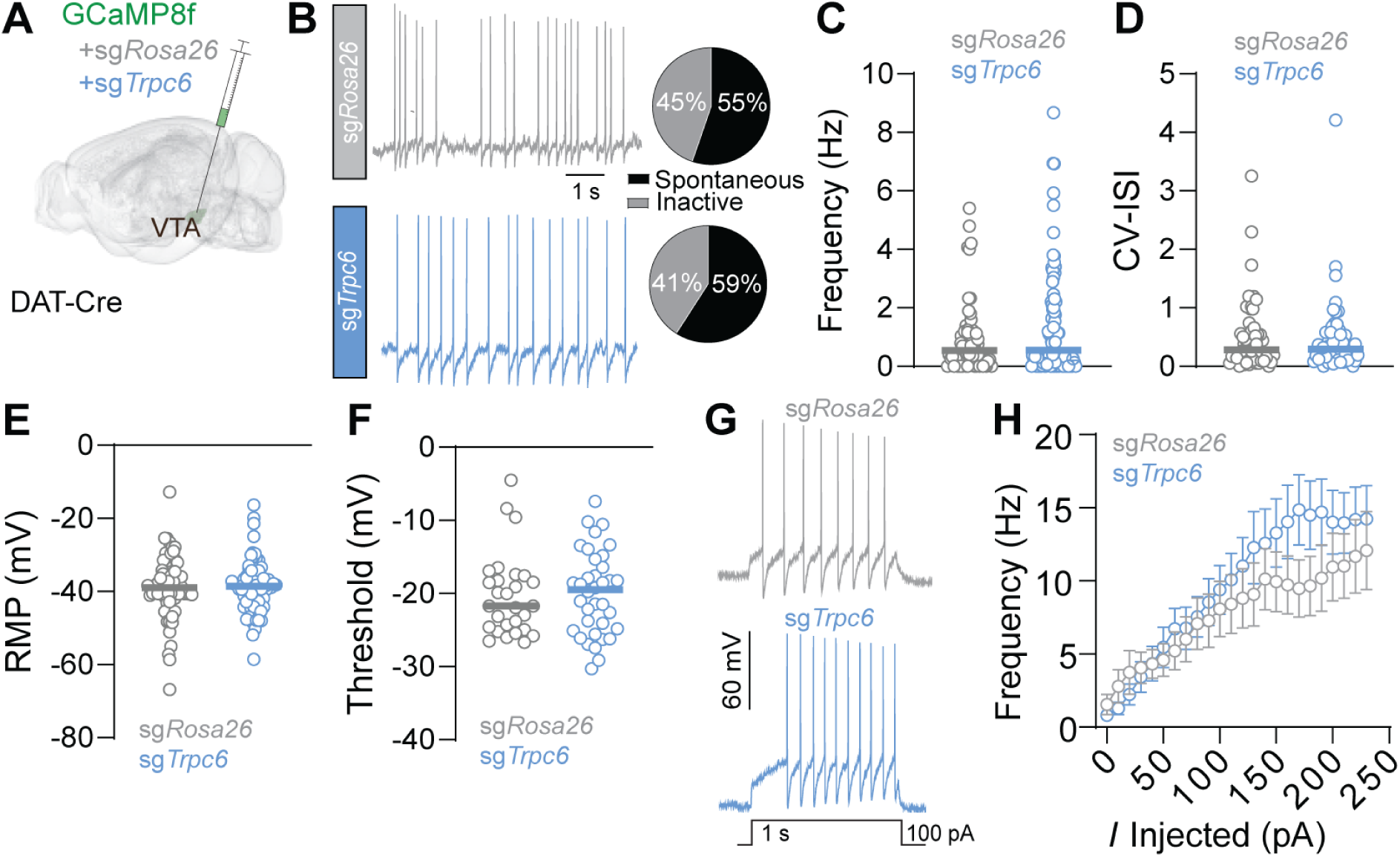
*Trpc6* mutagenesis does not alter spontaneous or evoked action potential firing in dopamine neurons. (A) Schematic illustrating viral injection into the VTA of DAT-Cre mice. (B) Representative traces of spontaneous action potential firing in sg*Rosa26* and sg*Trpc6* injected mice. The proportion of spontaneously active versus inactive cells did not differ significantly (N=5 sg*Rosa26* mice, n=76 cells and N=6 sg*Trpc6* mice, n=66 cells, Fisher’s exact test). (C) Spontaneous firing frequency in sg*Rosa26* and sg*Trpc6* injected mice did not differ significantly (N=5 sg*Rosa26* mice, n=65 cells and N=6 sg*Trpc6* mice, n=80 cells; D’Agostino & Pearson Test, Mann Whitney). (D) Coefficient of variation in the interspike interval (CV-ISI) in sg*Rosa26* and sg*Trpc6* injected mice did not differ significantly (N=5 sg*Rosa26* mice, n=51 cells and N=6 sg*Trpc6* mice, n=56 cells; D’Agostino & Pearson Test, Mann Whitney). (E) Resting membrane potential (RMP) in sg*Rosa26* and sg*Trpc6* injected mice did not differ significantly (N=5 sg*Rosa26* mice, n=65 cells and N=6 sg*Trpc6* mice, n=67 cells; D’Agostino & Pearson Test, Mann Whitney). (F) Threshold to fire action potential during ramp current injection in sg*Rosa26* and sg*Trpc6* injected mice did not differ significantly (N=8 sg*Rosa26* mice, n=28 cells and N=3 sg*Trpc6* mice, n=36 cells; D’Agostino & Pearson Test, Mann Whitney). (G) Example traces following current injection in sg*Rosa26* and sg*Trpc6* injected mice. (H) Spikes fired following current injection did not differ between sg*Rosa26* and sg*Trpc6* injected mice (N=13 sg*Rosa26* mice, n=15 cells and N=9 sg*Trpc6* mice, n=26 cells; Two-way repeated measures ANOVA, Bonferroni’s multiple comparisons). Data are presented as the mean ± S.E.M.

### TRPC6 channels regulate consummatory behavior and DA neuron activity in a homeostatic state-dependent manner

The neuropeptides NTS, Tac2, ghrelin, and angiotensin II are associated with energy balance and fluid/food consumption, and they signal through G_q/11_-coupled receptors. All of these neuropeptides have been shown to modulate DA neuron activity (Cone et al., 2014; Hsu et al., 2020; Naef et al., 2015; Palmiter, 2007), and NTS has been shown to regulate food reward-associated increase in calcium in VTA-DA neurons *in vivo* in hungry mice (Soden et al., 2023). Our *ex vivo* analysis of calcium signaling dynamics in VTA-DA neurons indicates that TRPC6 channels act directly downstream of stimulatory neuropeptide receptor signaling. Therefore, we asked whether TRPC6 channels influence *in vivo* calcium signaling dynamics of VTA-DA neurons during consummatory behavior in hungry or thirsty mice. To address this, we performed fiber photometry recordings in DAT-Cre mice injected with either AAV1-FLEX-sg*Rosa26* or AAV1-FLEX-sg*Trpc6*(1) and AAV1-FLEX-sg*Trpc6*(2), along with AAV1-FLEX-GCaMP6m into the VTA. An optic fiber was implanted over the injection site (Fig. 4A; Supplementary Fig. 1).

**Figure 4.**
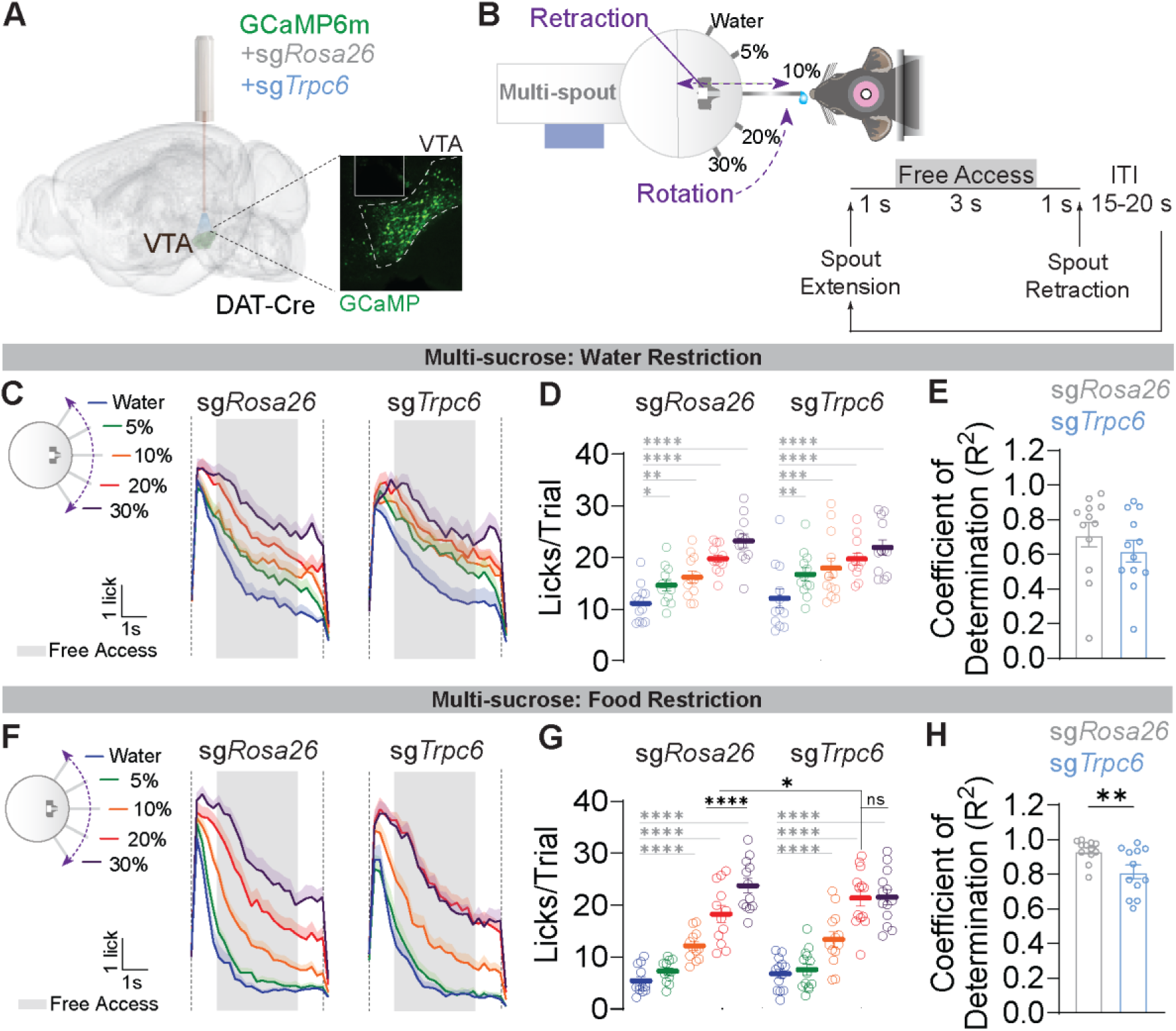
TRPC6 channels modulate licking behavior in a multi-sucrose consumption task under food restriction but not water restriction. (A) Schematic illustrating viral injections and optic fiber implants in the VTA of DAT-Cre mice. Inset is a representative histology image from immunohistochemistry validation of targeting. (B) Schematic of the OHRBETS system and trial structure that consists of a rotating and retracting five-spout apparatus. Head-restrained mice are presented with one spout at a time. Spout extension is followed by a 1 sec delay, 3 sec free access delivery of liquid, 1 sec delay, spout retraction, then a variable 15-20 sec inter-trial interval (ITI). (C-E) Multi-sucrose experiment under 90% water restriction. Lick rate graphs (C) and quantification of total licks per trial (D) for each spout identity (N=12 mice for both groups; Two-way repeated measures ANOVA, Tukey’s multiple comparisons, *P<0.05, **P<0.01, ***P<0.001, ****P<0.0001). (E) Regression coefficient analysis per animal (N=12 mice for both groups; Independent t-test, P=ns). (F-H) Multi-sucrose experiment under 85% food restriction. Lick rate graphs (F) and quantification of total licks per trial (G) for each spout identity (N=12 mice for both groups; Two-way repeated measures ANOVA, Tukey’s multiple comparisons, *P<0.05, ****P<0.0001). (H) Regression coefficient analysis per animal (N=12 mice for both groups; Independent t-test, **P<0.01). See Extended Data Table 1 for details on statistical tests. Data are presented as the mean ± S.E.M. averaged across three days of the experiment for C-H.

Consummatory licking behavior and photometry recordings were performed in head-restrained mice. Sucrose solutions of varying concentrations were delivered to water-or calorie-restricted mice using a multi-spout system previously described (Gordon-Fennell et al., 2023) (Fig. 4B). Briefly, mice were water restricted to 90% of baseline weight, head restrained and trained to lick five spouts for different concentrations of sucrose (water, 5%, 10%, 20%, and 30% sucrose) across five days. The spout identity was changed daily, and each spout was presented twelve times in a pseudorandom order within a daily session. The trial structure included a three second access period where mice could freely lick for liquid delivery (Fig. 4B). Following water restriction (WR), mice were given *ad libitum* access to water and then calorie restricted to 85% baseline body weight, and the multi-spout experiment was repeated as above.

Under WR, both groups exhibited similar licking behavior for all five solutions (Fig. 4C-D). Linear regression analysis of licks versus spout identity was not significantly different between groups (Fig. 4E). In contrast, under food restriction (FR), sg*Trpc6* mice reached maximal licking at 20% sucrose, whereas *sgRosa26* mice showed maximal licking at 30% sucrose (Fig. 4F-G). Linear regression analysis under FR revealed a significantly stronger relationship between licks and spout identity in sg*Rosa26* control mice relative to sg*Trpc6* mice (Fig. 4H). Both groups showed significantly more licking to water, 5%, and 10% sucrose under WR compared to FR and the effect of spout identity for WR versus FR was significantly greater in both groups (Supplementary Fig. 2C).

Analysis of calcium signals in VTA-DA neurons during the solution access period in the multi-spout assay under WR revealed nearly identical responses in sg*Rosa26* and sg*Trp6* mice (Fig. 5A-B) and the linear relationship between spout identity and calcium signals was similar (Fig. 5C). Under FR, however, sg*Rosa26* mice showed significantly higher calcium signals to 30% sucrose than sg*Trpc6* mice (Fig. 5D-E). While the linear regression analysis of area under the curve (AUC) during the solution access period versus spout identity was not significant between groups (Fig. 5F), within group comparisons revealed higher calcium signals to 30% sucrose compared to 20% sucrose in sg*Rosa26* mice that was not observed in sg*Trpc6* mice (Fig. 5E).

**Figure 5.**
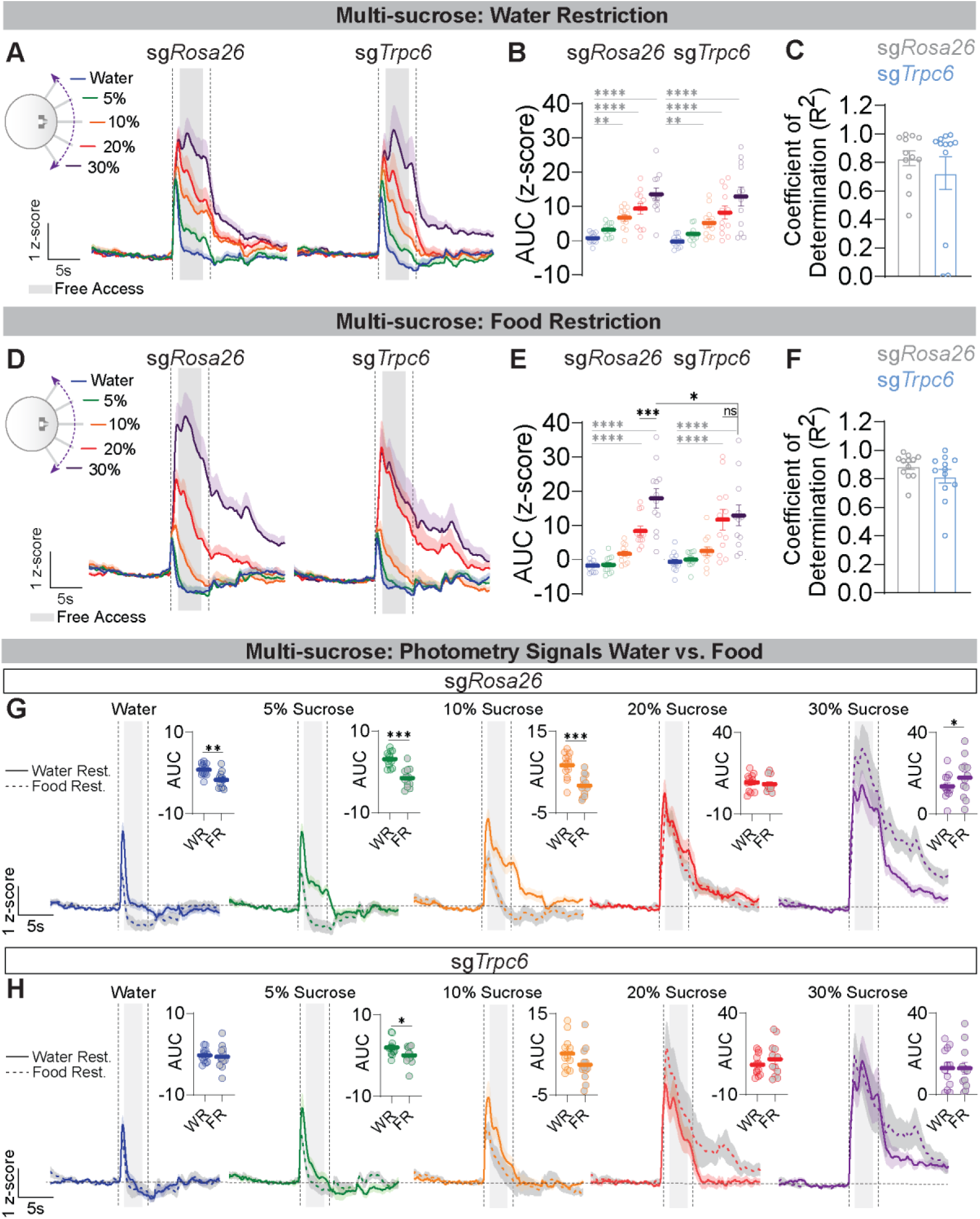
TRPC6 channels regulate *in vivo* VTA-DA calcium signals during a multi-sucrose consumption task under food restriction but not water restriction. (A-C) Multi-sucrose experiment under 90% water restriction. Z-scored fiber photometry signals aligned to spout extension (A) and AUC quantified during the 3 sec free access period (B) for each spout identity (N=12 mice for both groups; Two-way repeated measures ANOVA, Tukey’s multiple comparisons, **P<0.01, ****P<0.0001). (C) Regression coefficient analysis per animal (N=12 mice for both groups; Independent t-test, P=ns). (D-F) Multi-sucrose experiment under 85% food restriction. Z-scored fiber photometry signals aligned to spout extension (D) and AUC quantified during the 3 sec free access period (E) for each spout identity (N=12 mice for both groups; Two-way repeated measures ANOVA, Tukey’s multiple comparisons, *P<0.05, ***P<0.001, ****P<0.0001). (F) Regression coefficient analysis per animal (N=12 mice for both groups; Independent t-test, P=ns). Comparisons of z-scored fiber photometry signals and AUC quantifications from A, B, D, E between water restriction and food restriction experiments for each spout identity for sg*Rosa26* mice (G) and sg*Trpc6* mice (H) (N=12 mice for both groups; Paired samples t-test, *P<0.05, **P<0.01, ***P<0.001). See Extended Data Table 1 for details on statistical tests. Data are presented as the mean ± S.E.M. averaged across three days of the experiment for A-H.

Based on the differential effects of *Trpc6* mutagenesis on lick rates and calcium signals between WR and FR, we asked whether these effects were reflected as differences in the calcium signals within groups under these conditions. Comparison of calcium signals in sg*Rosa26* mice revealed significantly reduced responses to water, 5%, and 10% sucrose under WR compared to FR, and significantly greater responses to 30% sucrose under FR (Fig. 5G). Interestingly, sg*Rosa26* mice showed a dip in the calcium signal below baseline during the solution access period under FR, indicative of a reward prediction error (RPE)-like signal (Schultz et al., 1997). In contrast, sg*Trpc6* mice showed only modest but significantly greater calcium signals to 5% sucrose under WR compared to FR, and no difference under other conditions (Fig. 5H). Moreover, these animals did not show the same RPE-like signal under FR that we observed in sg*Rosa26* mice.

VTA-DA neurons have been shown to differentially encode homeostatic states under WR or FR, with cells showing increased activity in the post-consummatory phase following water consumption in thirsty mice (Grove et al., 2022). To address whether TRPC6 channels in VTA-DA neurons influence calcium signals post-consumption, we analyzed the AUC during the post-solution access period. We observed differential responses in sg*Rosa26* mice with higher post-consummatory activity at 10% sucrose with WR greater than FR and at 30% sucrose with FR greater than WR (Supplementary Fig. 3A). In sg*Trpc6* mice, we did observe a significant difference at 20% sucrose with FR greater than WR (Supplementary Fig. 3B). Additionally, similar to consummatory responses and calcium signals during the access period under FR, we observed significant differences between 20% and 30% sucrose in sg*Rosa26* mice during the post-consummatory period that was not significant in sg*Trpc6* mice (Supplementary Fig. 3A-B). We observed no differences between the groups in the linear relationship between post-consummatory calcium signals and spout identity (Supplementary Fig. 3C). Together, these findings demonstrate that TRPC6 channel signaling in DA neurons plays an important role in regulating consummatory responses and the differential encoding of solution content under varying homeostatic states, particularly during the solution access period.

## DISCUSSION

Our findings provide substantial evidence for the specific coupling of TRPC6 channels to neuropeptide receptor signaling pathways in VTA-DA neurons. We demonstrate selective modulation of calcium dynamics following neuropeptide receptor activation of Ntsr1 and Tacr3, whereas activation of other GPCRs (e.g. mGluR, mAChR) that also couple to G_q/11_ signaling do not engage TRPC6 channels. These GPCR signaling pathways involve PLC-mediated cleavage of PIP_2_ into IP_3_ and DAG. Despite the high likelihood that the same signaling cascade is utilized by both neuropeptide and neurotransmitter receptors, we observed highly selective coupling of TRPC6 channels to neuropeptide receptors. The robust calcium responses to DHPG and carbachol are consistent with previous reports of expression of mGluR5 and m5AchR in VTA-DA neurons (Merrill et al., 2015; Yeomans et al., 2001). This receptor-channel specificity may be attributed to the spatial proximity of TRPC channels relative to their activating GPCRs (Luo et al., 2025).

Indeed, scaffolding proteins like A-kinase anchoring proteins (AKAPs) spatially constrain GPCR and ion channel signaling (Wong and Scott, 2004). The proximity between mAChRs and TRPV4 channels in endothelial cells is regulated by AKAP79/150, and the interaction between TRPV4 and AKAP79/150 is critical for carbachol-induced activation of the channel (Sonkusare et al., 2014). Little is known about interactions between AKAPs and TRPC6 channels; however, AKAP79/150 protein has been detected in VTA-DA neurons (Dacher et al., 2013). Future experiments to determine the signaling complexes involved in the control of TRPC6 activation by neuropeptide receptors will be highly informative.

The functional implications of neuropeptide receptor-TRPC6 channel selective coupling are supported by our *in vivo* data. Our results demonstrate that *Trpc6* mutagenesis directly impacts VTA-DA neuron calcium signaling dynamics associated with consummatory behavior and caloric content. Specifically, we show that TRPC6 channels play an important role in encoding sucrose concentration in a manner that is consistent with reward value encoding and modulating consummatory behavior in response to specific homeostatic demands (hunger versus thirst). Between group comparisons of calcium signals during the multi-sucrose task under food restriction reveal that mutagenesis of *Trpc6* impairs the scaling of VTA-DA neuron calcium signals and subsequent behavioral responses to high-value sucrose rewards.

Within-group comparisons reveal a more complex relationship between TRPC6 channel signaling and scalar encoding of value. Both sg*Rosa26* control and sg*Trpc6* mutants showed increased calcium signals in DA neurons to increasing sucrose concentrations under both WR and FR. However, the dynamics of these signals across sucrose concentrations differed markedly between WR and FR in sg*Rosa26* control mice but were largely similar in sg*Trpc6* mutant mice. Results observed in *sgRosa26* control mice are consistent with findings that DA neurons differentially encode information relating to hydration and caloric intake under thirst and hunger conditions (Grove et al., 2022). These results suggest that TRPC6 channel activation, likely downstream of specific neuropeptide receptors, functions in a state-dependent manner.

Our *in vivo* results also demonstrate a broader range of value encoding in sg*Rosa26* control mice—but not *sgTrpc6* mice—under FR compared to WR, suggesting differential activation of neuropeptide receptors that are coupled to TRPC6 channels. The neuropeptides that recruit activation of TRPC6 channels in hungry mice are not yet known, but candidates include NTS, which has been shown to regulate feeding (Gazit Shimoni et al., 2025; Ramirez-Virella and Leinninger, 2021), and the gut-derived peptide ghrelin that is also G_q/11_ coupled and enhances dopamine neuron activity (Abizaid et al., 2006). In contrast, DA responses to thirst are regulated by renin/angiotensin signaling, but this effect is thought to be mediated by angiotensin II receptor signaling outside the VTA (Hsu et al., 2020).

In conclusion, our findings support previous work illustrating how neuropeptide receptor activation of TRPC channels tightly regulates neuronal excitability and synchronized activity (Kelly and Wagner, 2024). We demonstrate a novel mechanism by which TRPC6 channels are coupled to neuropeptidergic signaling pathways to regulate VTA-DA neuron activity and behavior under specific homeostatic demands. Importantly, our electrophysiological data confirm that the differences we observe in calcium signaling after *Trpc6* mutagenesis are not due to changes in intrinsic or evoked excitability. The lack of effect on baseline neuronal properties underscores the therapeutic potential of TRPC6 channels, as interventions targeting these channels could modulate pathological signaling without disrupting essential neuronal function. These results provide evidence for TRPC6 channels as a potential therapeutic target for a variety of disorders related to imbalances in consummatory and reward-related behaviors.

## MATERIALS AND METHODS

### Experimental Subject Details

Male and female mice, housed on a 12-h light/dark cycle, were used in all experiments performed during the light phase. All procedures were approved and performed under the guidelines of the Institutional Animal Care and Use Committee at the University of Washington (PROTO201600703 – 4249-01). Mice between the ages of 2–6 months were used for all experiments. DAT-Cre mice (B6.SJL-*Slc6a3^tm1.1(cre)Bkmn^*/J) were purchased from The Jackson Laboratory and bred in-house.

### Viruses

All AAV1 viruses were produced in-house with titers of 1×10^12^ to 3×10^12^ particles per mL as described (Gore et al., 2013).

### Design of CRISPR Constructs

Primers used for cloning sg*Trpc6*(1) into AAV1-FLEX-SaCas9-U6 were: forward CACCGAGCCTTTAGAGAGCCGCCCCT and reverse AAACAGGGGCGGCTCTCTAAAGGCTC. Primers used for cloning sg*Trpc6*(2) into AAV1-FLEX-SaCas9-U6 were: forward CACCGTCCTACTACATTGGCGCAAAA and reverse AAACTTTTGCGCCAATGTAGTAGGAC. The control CRISPR virus used was AAV1-FLEX-SaCas9-U6-sg*Rosa26,* and single-guide RNAs targeting *Trpc6* were designed as previously published (Hunker et al., 2020; Hunker and Zweifel, 2020). All CRISPR viruses contained a hemagglutinin (HA) tag used to confirm virus expression using immunohistochemistry.

### Validation of Mutagenesis

Validation of mutagenesis was performed using fluorescence-activated cell sorting, whole-genome amplification and sequencing as described (Hunker et al., 2020; Hunker and Zweifel, 2020). In brief, three DAT-Cre mice were injected in the VTA with AAV1-FLEX-SaCas9-U6-sg*Trpc6*(1) and AAV1-FLEX-SaCas9-U6-sg*Trpc6*(2) along with AAV-FLEX-EGFP-KASH to label nuclei for sorting. Four weeks following injection tissue punches of the VTA were collected and GFP-positive nuclei were isolated using fluorescence-activated cell sorting. Whole-genome amplification (REPLI-g, Qiagen) was performed according to the manufacturer’s instructions, followed by targeted sequencing of a 200–300 bp region surrounding the intended cut site. Tracking of indels by decomposition (TIDE) analysis (Brinkman et al., 2014) was performed to compare sequence chromatograms from GFP-negative and GFP-positive samples to estimate mutation frequency.

### Surgery

Mice were anesthetized with isoflurane (1.5–4%) before and during viral injections and fiber implantations. Mice recovered for at least 4 weeks prior to experimentation. For slice electrophysiology, mice were injected at approximately 5–6 weeks of age. For all other experiments, mice were injected at 8–12 weeks of age. VTA coordinates were M–L: ±0.5, A–P: −3.25, D–V: −4.4. Values are in mm, relative to bregma. A–P values were adjusted for bregma– lambda distance using a correction factor of 4.21 mm. For z-values the syringe was lowered 0.5 mm past the indicated depth and raised up at the start of the injection. Injection volume was 500 nL at a rate of 250 nL/min. All injections were bilateral.

For electrophysiology and *ex vivo* calcium imaging experiments, all mice were injected with AAV1-FLEX-GCaMP8f at a 1:4 dilution. The diluent was the CRISPR virus: AAV1-FLEX-sg*Rosa26* (control) or AAV1-FLEX-sg*Trpc6*(1) and AAV1-FLEX-sg*Trpc6*(2) (experimental group). For fiber photometry experiments, all mice were injected with AAV1-FLEX-GCaMP6m at a 1:6 dilution. The diluent was the CRISPR virus: AAV1-FLEX-sg*Rosa26* (control) or AAV1-FLEX-sg*Trpc6*(1) and AAV1-FLEX-sg*Trpc6*(2) (experimental group).

Fiber-optic cannulas for photometry (400 µm fiber, 0.66 NA, 1.25 mm ferrule) were purchased from Doric and implanted in the VTA at a depth of −4.2 mm from bregma. Implantations were unilateral and implanted in the left hemisphere.

### *Ex Vivo* Slice Physiology

Horizontal brain slices (200 µm) were prepared in artificial cerebrospinal fluid (ACSF) solution 32 °C containing (in mM): 126 NaCl, 2.5 KCl, 1.2 MgCl_2_, 2.4 CaCl_2_, 1.2 NaH_2_PO_4_, 21.4 NaHCO_3_, 11.1 D-glucose and 10 µM MK-801 (to prevent NMDA-mediated excitotoxicity). Slices recovered in ACSF (in mM: 126 NaCl, 2.5 KCl, 1.2 NaH_2_PO4, 1.2 MgCl_2_ 11 d-glucose, 18 NaHCO_3_, 2.4 CaCl_2_) at 32°C containing 10 µM MK-801 for ≥ 30 minutes. All solutions were continually bubbled with O_2_ and CO_2_.

Slices were hemisected along the midline, and one half was transferred to a recording chamber under continuous perfusion (∼2 ml min^−1^) of oxygenated ACSF at 32 °C while the other was placed back into recovery. Recordings began in a whole-cell voltage clamp configuration with glass microelectrodes (World Precision Instruments) with a resistance of 2-3 MΩ. All electrophysiology experiments used an internal solution containing (in mM): 130 K-gluconate, 10 HEPES, 5 NaCl, 1 EGTA, 5 Mg-ATP, 0.5 Na-GTP, pH 7.3, 280 mOsm. Once a stable recording was achieved at-60 mV, cells were switched to current clamp and the spontaneous activity that followed for the next 70 seconds was recorded. Cells were then given escalating current steps (Δ = +10 pA) from-60 pA to +230 pA. Finally, cells were given a ramp protocol from 0 to 200 pA over 2 seconds. Basal electrical properties were measured and monitored following break-in using the average of eight 5 mV pulses (10 ms pulse, sampled at 10 kHz). Cells with a series resistance ≥16 MΩ, or with a change in series resistance ≥50% during the recording period were excluded from analysis. Whole-cell recordings were made using an Axonpatch 700B amplifier (Molecular Devices) and Clampex software.

### Electrophysiology Data Analysis

Electrophysiological data were acquired using Clampex software and analyzed post hoc with custom Python scripts. Recordings consisted of single or multiple sweeps loaded with the Neo library and AxonIO module. Spontaneous action potentials were detected by calculating the first derivative of voltage (dV/dt) and applying a threshold of ≥15 mV/ms. To quantify spontaneous firing rates, a sliding window analysis (10-second duration) was performed to identify the highest frequency of spontaneous action potentials within a 70-second recording period.

Resting membrane potential was estimated by identifying interspike intervals of at least 200 ms and calculating the average voltage over a 50-point window within these quiescent periods. This approach ensured that measurements reflected the membrane’s most hyperpolarized state and were not influenced by pre-spike depolarizations or post-spike afterpotentials.

Spike width was determined by identifying action potential boundaries: the spike onset was defined as the first point before the detected peak where dV/dt ≤ 0, and repolarization was identified as the point following the peak where dV/dt returned to ≥ 0. Spike width was subsequently measured between these two boundary points. The coefficient of variability (CV) of inter-spike intervals (ISI) was calculated to assess firing regularity.

Additionally, rheobase currents were determined from ramp current injections by calculating the minimum current required to elicit an action potential using robust spike validation criteria. Spikes during ramp protocols were validated based on sustained threshold crossing of dV/dt (≥ 0.1× peak dV/dt for ≥ 0.5 ms) and confirmed depolarization above 0 mV. The spike threshold voltage was defined as the membrane potential at the first point of dV/dt threshold crossing that led to a full action potential and was used to quantify the voltage level at which each neuron initiated firing.

### *Ex Vivo* Slice Calcium Imaging

Horizontal brain slices (200 µm) were prepared in an ice slush solution containing (in mM): 92 *N*-methyl-d-glucamine, 2.5 KCl, 1.25 NaH_2_PO4, 30 NaHCO_3_, 20 HEPES, 25 glucose, 2 thiourea, 5 sodium ascorbate, 3 sodium pyruvate, 0.5 CaCl_2_, 10 MgSO_4_, pH 7.3–7.4 (Ting, 2014). Slices recovered for 12 minutes in the same solution at 32 °C and then were transferred to a room temperature solution including (in mM): 92 NaCl, 2.5 KCl, 1.25 NaH_2_PO_4_, 30 NaHCO_3_, 20 HEPES, 25 glucose, 2 thiourea, 5 sodium ascorbate, 3 sodium pyruvate, 2 CaCl_2_, 2 MgSO_4_. Slices recovered for an additional 45 minutes before recordings were made in ACSF at 32 °C that was continually perfused over slices at a rate of ∼2 ml min^−1^ and contained (in mM): 126 NaCl, 2.5 KCl, 1.2 NaH_2_PO4, 1.2 MgCl_2_ 11 d-glucose, 18 NaHCO_3_, 2.4 CaCl_2_. All solutions were continually bubbled with O_2_ and CO_2_.

For bath application experiments, a 3-minute baseline period was recorded. Senktide (500 nM) was applied for two minutes, then a 10-minute washout period followed. For puffing experiments, ∼10 MΩ electrodes were filled with either GSK 1702934A (1 µM and 300 nM), (S)-3,5-Dihydroxyphenylglycine (30 µM), neurotensin (1 µM), senktide (500 nM), or carbachol (30 µM) diluted with an internal solution (in mM): 126 NaCl, 2.5 KCl, 1.2 NaH_2_PO_4_, 1.2 MgCl_2_ 11 d-glucose, 18 NaHCO_3_, 2.4 CaCl_2_.

The puffing pipette was situated adjacent to the cell of interest, and only one cell was analyzed per puff. Only cells that exhibited a visual mechanical deflection in response to the puff were subsequently analyzed. Importantly, while this deflection was reflected by a sharp negative trough in the calcium signal, the calcium signal immediately rebounded. Cells that exhibited a deflection and an increase in calcium signal were classified as responders; cells that exhibited the deflection without increase in calcium fluorescence were categorized as nonresponders (see Fig. 1H inset for an example). Images were acquired using a high-speed camera (Hamamatsu OrcaFlash 4.0LT) and Metafluor software.

### *Ex Vivo* Slice Calcium Imaging Data Analysis

ΔF/F was calculated from the 3-minute baseline period using ImageJ and Inscopix software. Peak fluorescence and area under the curve were compared between the two groups using a Student’s t-test.

### Fiber Photometry

Mice were connected to an imaging patch cord (Doric Lenses) and head-fixed in the multi-spout apparatus. The imaging patch cord was photobleached for 10 minutes prior to recording. Recordings were made using an RZ5 BioAmp Processor and Synapse software (Tucker Davis Technologies). A 465 nm LED (531-Hz, sinusoidal, Doric Lenses) was used to excite GCaMP6m. LED intensity was measured at the tip of the optic fiber prior to each recording session and set to 30–40 µW. GCaMP6m fluorescence (525 ± 25 nm) was returned through the same patch cord, bandpass filtered, and recorded by the RZ5 at a sampling rate of 1,017.25 Hz. Spout extension events were synced to the photometry recording via TTL delivery for offline analysis.

### Fiber Photometry Data Analysis

The 465 nm signal was preprocessed as previously described (Simpson et al., 2024). In brief, the signal was down sampled 100x using a moving window mean and corrected for photobleaching using a double exponential curve fit and subtraction (Simpson et al., 2024). A custom Python package adapted from the Tucker Davis code was used to extract and analyze the GCaMP signal surrounding each event. A 30-second window was extracted surrounding each spout extension (10 seconds prior and 20 seconds following). Five seconds to one second prior to spout extension was used as a baseline to calculate the z-score. Z-scores for each unique spout identity were averaged for each mouse.

### Multi-spout Apparatus

Mice were head-fixed in an OHRBETS (Open-Source Head-fixed Rodent Behavioral Experimental Training System) as described (Gordon-Fennell et al., 2023). Briefly, OHRBETS was constructed from 3D printed components and Arduino hardware. Mice were trained to lick for solutions from multiple rotating and retracting spouts. Under water restriction, mouse weights were held consistent at 90% baseline. Under food restriction, mouse weights were held consistent at 85% of baseline weight. Multi-spout experiments were completed in four cohorts of mice.

### Habituation to Head Fixation and Free-access Lick Training

Mice were water restricted to 90% baseline weight. Mice were habituated to head fixation and trained to lick for 10% sucrose in a single 10-minute session. Mice were scruffed and gently had their rear end and hind paws placed in a 50 mL conical during head fixation. During this session, mice were given free access to 10% sucrose. Free access was approximated using closed-loop delivery of a pulse of fluid (∼1.5 µL) each time the mouse licked the spout. The spout was positioned ∼2–3 mm in front of the mouse’s mouth where it remained throughout the session (Gordon-Fennell et al., 2023).

### Retractable Spout Training

Mice were water restricted to 90% of baseline weight. Retractable spout training consisted of two daily sessions of 60 trials using one spout. The trial structure was as follows: spout extension, one second delay with no liquid available, 3 seconds of free access of 10% sucrose as described above, a one second delay with no liquid delivery, spout retraction. The inter-trial interval was a variable 15-20 seconds.

### Multi-spout Brief Access Sucrose Experiment

Mice were either food-restricted or water-restricted. Mice underwent five consecutive days of multi-spout brief access for five concentrations of sucrose (0%, 5%, 10%, 20%, 30%) across 60 trials. The trial structure was as follows: spout extension, one second delay with no liquid available, 3 seconds of free access of 10% sucrose, a one second delay with no liquid delivery, spout retraction. The inter-trial interval was a variable 15-20 seconds. Each spout contained a different concentration of sucrose that remained the same throughout the daily session. The solutions in the spouts, as well as the order of the spout presentations, were pseudorandomly assigned each day. Fiber photometry signals were recorded during the third, fourth and fifth days of the experiment. Licking and photometry data are presented as the average of these three days for each mouse.

### Immunohistochemistry

Mice were deeply anesthetized using Beuthanasia and transcardially perfused with phosphate-buffered saline (PBS), followed by 4% paraformaldehyde (PFA) in PBS. Brains were placed in 4% PFA overnight, then transferred to 30% sucrose in PBS solution at 4°C for at least 48 hours. Brains were then sectioned into 45μm sections using a cryostat and placed in PBS at 4°C. Brain sections were then stained to validate virus expression. Free-floating sections were placed in a blocking buffer (3% normal donkey serum and 0.3% Triton X-100 in PBS) for 30 minutes. For enhancement of GCaMP6m and HA tag, sections were incubated in primary antibodies (Chicken-GFP, 1:6000 dilution, ABCAM, Rabbit-HA, 1:1000) overnight at 4°C. Sections were then placed in PBS for three 10-minute washes and incubated in secondary antibody (1:250 dilution Alexa Fluor 488 AffiniPure Donkey Anti-Chicken or Cy3 Donkey Anti-Rabbit, Jackson ImmunoResearch) for 45 minutes at room temperature. Then, following three 10-minute PBS washes, sections were mounted using a mounting medium (DAPI Fluoromount-G, Southern Biotech) and cover-slipped. Images were collected using a Keyence BZ-X710 fluorescent microscope and analyzed using ImageJ software.

### Statistics and Reproducibility

All data were analyzed for statistical significance using Prism software (GraphPad Prism 9). The Geisser–Greenhouse correction was used to correct for unequal variability of differences in repeated-measures ANOVA tests. No statistical methods were used to predetermine sample sizes. All behavioral assays were repeated in a minimum of two cohorts with similar replication of results. Littermates were randomly assigned to experimental groups and mice were tested in random order. Mice with missed viral injections or significant viral spread outside the targeted region were excluded from analyses.

## Acknowledgments

This work was supported by National Institutes of Health grants R01MH104450 (LSZ), R01DA044315 (LSZ), and F31DA058381 (MXB). We thank Dr. Selena Schattauer for assistance with viral production and the FACS experiment, as well as the University of Washington Center of Excellence in Opioid Addiction Research/Molecular Genetics Resource Core (P30DA048736). We thank Dr. Marta Soden and Mary Loveless for assistance with the multi-spout apparatus. We also thank members of the Zweifel, Soden, and Palmiter labs for their thoughtful discussions.

## Author contributions

M.X.B. and L.S.Z. conceptualized the project and experimental design for all experiments. M.X.B. acquired the puff slice calcium imaging and fiber photometry datasets.

M.T. collected the bath application slice calcium imaging dataset. O.K. collected and analyzed the electrophysiology datasets. M.X.B. and D.T.M. wrote Python scripts and analyzed all the datasets.

A.F. and L.S.Z. assisted with additional data analyses. S.J. aided with construction of the multi-spout apparatus and provided thoughtful discussion. M.X.B. and L.S.Z wrote the manuscript. All authors reviewed the manuscript.

## Data Availability

All data associated with this study will be made available by the corresponding author upon reasonable request.

## Code Availability

Code for fiber photometry analysis was derived from a publicly available source (Tucker Davis Technologies) and is available through GitHub (https://github.com/molliebernstein).

## References

1. Abizaid, A., Liu, Z.W., Andrews, Z.B., Shanabrough, M., Borok, E., Elsworth, J.D., Roth, R.H., Sleeman, M.W., Picciotto, M.R., Tschop, M.H., et al. (2006). Ghrelin modulates the activity and synaptic input organization of midbrain dopamine neurons while promoting appetite. J Clin Invest 116, 3229–3239.

2. Brinkman, E.K., Chen, T., Amendola, M., and van Steensel, B. (2014). Easy quantitative assessment of genome editing by sequence trace decomposition. Nucleic Acids Res 42, e168.

3. Chen, L.W., Guan, Z.L., and Ding, Y.Q. (1998). Mesencephalic dopaminergic neurons expressing neuromedin K receptor (NK3): a double immunocytochemical study in the rat. Brain Res 780, 150–154.

4. Chung, A.S., Miller, S.M., Sun, Y., Xu, X., and Zweifel, L.S. (2017). Sexual congruency in the connectome and translatome of VTA dopamine neurons. Sci Rep 7, 11120.

5. Clapham, D.E. (1995). Calcium signaling. Cell 80, 259–268.

6. Cone, J.J., McCutcheon, J.E., and Roitman, M.F. (2014). Ghrelin acts as an interface between physiological state and phasic dopamine signaling. J Neurosci 34, 4905–4913.

7. Dacher, M., Gouty, S., Dash, S., Cox, B.M., and Nugent, F.S. (2013). A-kinase anchoring protein-calcineurin signaling in long-term depression of GABAergic synapses. J Neurosci 33, 2650–2660.

8. Forman, C.J., Tomes, H., Mbobo, B., Burman, R.J., Jacobs, M., Baden, T., and Raimondo, J.V. (2017). Openspritzer: an open hardware pressure ejection system for reliably delivering picolitre volumes. Sci Rep 7, 2188.

9. Gazit Shimoni, N., Tose, A.J., Seng, C., Jin, Y., Lukacsovich, T., Yang, H., Verharen, J.P.H., Liu, C., Tanios, M., Hu, E., et al. (2025). Changes in neurotensin signalling drive hedonic devaluation in obesity. Nature 641, 1238–1247.

10. Gordon-Fennell, A., Barbakh, J.M., Utley, M.T., Singh, S., Bazzino, P., Gowrishankar, R., Bruchas, M.R., Roitman, M.F., and Stuber, G.D. (2023). An open-source platform for head-fixed operant and consummatory behavior. Elife 12.

11. Gore, B.B., Soden, M.E., and Zweifel, L.S. (2013). Manipulating gene expression in projection-specific neuronal populations using combinatorial viral approaches. Curr Protoc Neurosci 65, 4 35 31-20.

12. Grove, J.C.R., Gray, L.A., La Santa Medina, N., Sivakumar, N., Ahn, J.S., Corpuz, T.V., Berke, J.D., Kreitzer, A.C., and Knight, Z.A. (2022). Dopamine subsystems that track internal states. Nature 608, 374–380.

13. Hsu, T.M., Bazzino, P., Hurh, S.J., Konanur, V.R., Roitman, J.D., and Roitman, M.F. (2020). Thirst recruits phasic dopamine signaling through subfornical organ neurons. Proc Natl Acad Sci U S A 117, 30744–30754.

14. Hunker, A.C., Soden, M.E., Krayushkina, D., Heymann, G., Awatramani, R., and Zweifel, L.S. (2020). Conditional Single Vector CRISPR/SaCas9 Viruses for Efficient Mutagenesis in the Adult Mouse Nervous System. Cell Rep 30, 4303–4316 e4306.

15. Hunker, A.C., and Zweifel, L.S. (2020). Protocol to Design, Clone, and Validate sgRNAs for In Vivo Reverse Genetic Studies. STAR Protoc 1.

16. Kaczmarek, L.K. (1987). The role of protein kinase C in the regulation of ion channels and neurotransmitter release. Trends in Neurosciences 10, 30–34.

17. Kelly, M.J., Qiu, J., and Ronnekleiv, O.K. (2018). TRPCing around the hypothalamus. Front Neuroendocrinol 51, 116–124.

18. Kelly, M.J., and Wagner, E.J. (2024). Canonical transient receptor potential channels and hypothalamic control of homeostatic functions. J Neuroendocrinol 36, e13392.

19. Khan, R., Laumet, G., and Leinninger, G.M. (2024). Hungry for relief: Potential for neurotensin to address comorbid obesity and pain. Appetite 200, 107540.

20. Luo, Y., Sun, L., and Peng, Y. (2025). The structural basis of the G protein-coupled receptor and ion channel axis. Curr Res Struct Biol 9, 100165.

21. Mathes, C., and Thompson, S.H. (1994). Calcium current activated by muscarinic receptors and thapsigargin in neuronal cells. J Gen Physiol 104, 107–121.

22. Merrill, C.B., Friend, L.N., Newton, S.T., Hopkins, Z.H., and Edwards, J.G. (2015). Ventral tegmental area dopamine and GABA neurons: Physiological properties and expression of mRNA for endocannabinoid biosynthetic elements. Sci Rep 5, 16176.

23. Mietlicki-Baase, E.G., Santollo, J., and Daniels, D. (2021). Fluid intake, what’s dopamine got to do with it? Physiol Behav 236, 113418.

24. Naef, L., Pitman, K.A., and Borgland, S.L. (2015). Mesolimbic dopamine and its neuromodulators in obesity and binge eating. CNS Spectr 20, 574–583.

25. Nalivaiko, E., Michaud, J.C., Soubrie, P., Le Fur, G., and Feltz, P. (1997). Tachykinin neurokinin-1 and neurokinin-3 receptor-mediated responses in guinea-pig substantia nigra: an in vitro electrophysiological study. Neuroscience 78, 745–757.

26. Palmiter, R.D. (2007). Is dopamine a physiologically relevant mediator of feeding behavior? Trends Neurosci 30, 375–381.

27. Qiu, J., Nestor, C.C., Zhang, C., Padilla, S.L., Palmiter, R.D., Kelly, M.J., and Ronnekleiv, O.K. (2016). High-frequency stimulation-induced peptide release synchronizes arcuate kisspeptin neurons and excites GnRH neurons. Elife 5.

28. Qiu, J., Stincic, T.L., Bosch, M.A., Connors, A.M., Kaech Petrie, S., Ronnekleiv, O.K., and Kelly, M.J. (2021). Deletion of Stim1 in Hypothalamic Arcuate Nucleus Kiss1 Neurons Potentiates Synchronous GCaMP Activity and Protects against Diet-Induced Obesity. J Neurosci 41, 9688–9701.

29. Ramirez-Virella, J., and Leinninger, G.M. (2021). The Role of Central Neurotensin in Regulating Feeding and Body Weight. Endocrinology 162.

30. Schultz, W., Dayan, P., and Montague, P.R. (1997). A neural substrate of prediction and reward. Science 275, 1593–1599.

31. Seutin, V., Massotte, L., and Dresse, A. (1989). Electrophysiological effects of neurotensin on dopaminergic neurones of the ventral tegmental area of the rat in vitro. Neuropharmacology 28, 949–954.

32. Shin, K.C., Ali, G., Ali Moussa, H.Y., Gupta, V., de la Fuente, A., Kim, H.G., Stanton, L.W., and Park, Y. (2023). Deletion of TRPC6, an Autism Risk Gene, Induces Hyperexcitability in Cortical Neurons Derived from Human Pluripotent Stem Cells. Mol Neurobiol 60, 7297–7308.

33. Simpson, E.H., Akam, T., Patriarchi, T., Blanco-Pozo, M., Burgeno, L.M., Mohebi, A., Cragg, S.J., and Walton, M.E. (2024). Lights, fiber, action! A primer on in vivo fiber photometry. Neuron 112, 718–739.

34. Soden, M.E., Yee, J.X., and Zweifel, L.S. (2023). Circuit coordination of opposing neuropeptide and neurotransmitter signals. Nature 619, 332–337.

35. Sonkusare, S.K., Dalsgaard, T., Bonev, A.D., Hill-Eubanks, D.C., Kotlikoff, M.I., Scott, J.D., Santana, L.F., and Nelson, M.T. (2014). AKAP150-dependent cooperative TRPV4 channel gating is central to endothelium-dependent vasodilation and is disrupted in hypertension. Sci Signal 7, ra66.

36. Stuhrman, K., and Roseberry, A.G. (2015). Neurotensin inhibits both dopamine-and GABA-mediated inhibition of ventral tegmental area dopamine neurons. J Neurophysiol 114, 1734–1745.

37. Suh, B.C., and Hille, B. (2002). Recovery from muscarinic modulation of M current channels requires phosphatidylinositol 4,5-bisphosphate synthesis. Neuron 35, 507–520.

38. Wang, H., Cheng, X., Tian, J., Xiao, Y., Tian, T., Xu, F., Hong, X., and Zhu, M.X. (2020). TRPC channels: Structure, function, regulation and recent advances in small molecular probes. Pharmacol Ther 209, 107497.

39. Wang, J., Su, M., Zhang, D., Zhang, L., Niu, C., Li, C., You, S., Sang, Y., Zhang, Y., Du, X., et al. (2024). The cation channel mechanisms of subthreshold inward depolarizing currents in the mice VTA dopaminergic neurons and their roles in the chronic-stress-induced depression-like behavior. Elife 12.

40. Wang, Q., Wang, D., Shibata, S., Ji, T., Zhang, L., Zhang, R., Yang, H., Ma, L., and Jiao, J. (2019). Group I metabotropic glutamate receptor activation induces TRPC6-dependent calcium influx and RhoA activation in cultured human kidney podocytes. Biochem Biophys Res Commun 511, 374–380.

41. Wise, R.A. (2004). Dopamine, learning and motivation. Nat Rev Neurosci 5, 483–494.

42. Wong, W., and Scott, J.D. (2004). AKAP signalling complexes: focal points in space and time. Nat Rev Mol Cell Biol 5, 959–970.

43. Yeomans, J., Forster, G., and Blaha, C. (2001). M5 muscarinic receptors are needed for slow activation of dopamine neurons and for rewarding brain stimulation. Life Sci 68, 2449–2456.

44. Zhang, L., Guo, F., Kim, J.Y., and Saffen, D. (2006). Muscarinic acetylcholine receptors activate TRPC6 channels in PC12D cells via Ca2+ store-independent mechanisms. J Biochem 139, 459–470.

45. Zhang, M., Ma, Y., Ye, X., Zhang, N., Pan, L., and Wang, B. (2023). TRP (transient receptor potential) ion channel family: structures, biological functions and therapeutic interventions for diseases. Signal Transduct Target Ther 8, 261.

