## Supplementary Material for "A neuropeptide-specific signaling pathway for state-dependent regulation of the mesolimbic dopamine system"

**Supplemental Figures and Legends**

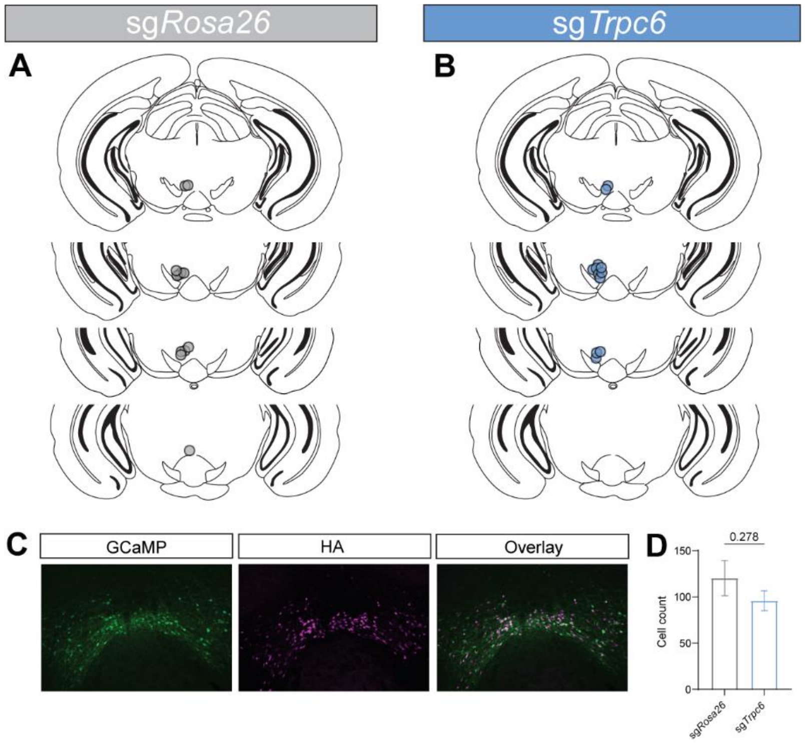

**Supplemental Figure 1.** Histological validation for fiber photometry mice. Optic fiber placements in the VTA for sg*Rosa26* mice (A) and sg*Trpc6* mice (B) (N=12 mice for both groups). (C) Representative immunohistochemistry images showing co-expression of GCaMP and hemagglutinin (HA) (CRISPR virus expression). (D) Cell counts for GCaMP-positive cells from fiber photometry mice in A and B (N=12 mice for both groups; Independent t-test, P=ns).

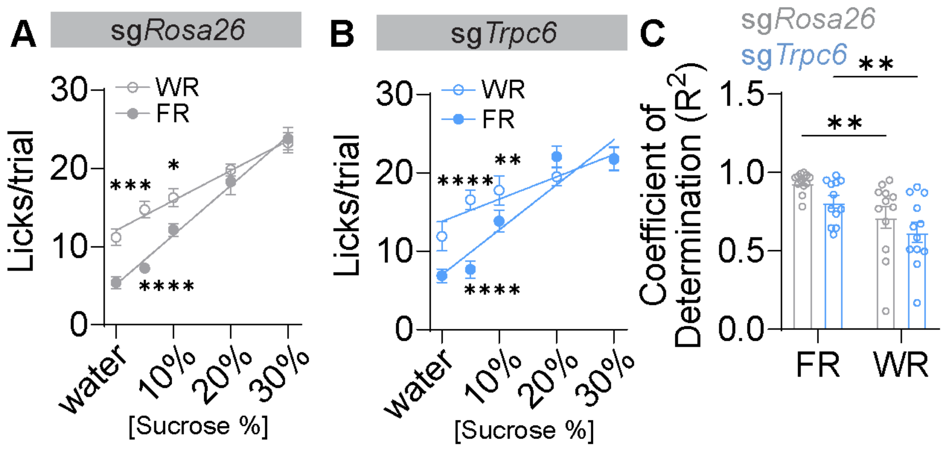

**Supplemental Figure 2.** Water restriction elicits higher licking responses at lower but not higher sucrose concentrations than food restriction in both sg*Rosa26* and sg*Trpc6* mice. (A-B) Licking responses for water and increasing sucrose in sg*Rosa26* (A) and sg*Trpc6* (B) mice. Two-way repeated measures ANOVA within subject comparison (*P<0.05, **P<0.01, ***P<0.001, and ****P<0.0001, Sidak post hoc comparison). (C) Linear regression analysis for individual mice reveals significant differences within sg*Rosa26* under food restriction (FR) versus water restriction (WR) and within sg*Trpc6* mice under FR versus WR (Two-way repeated measures ANOVA; **P<0.01, ***P<0.001, and ****P<0.0001, Sidak post hoc comparison). Data are presented as mean ± S.E.M.

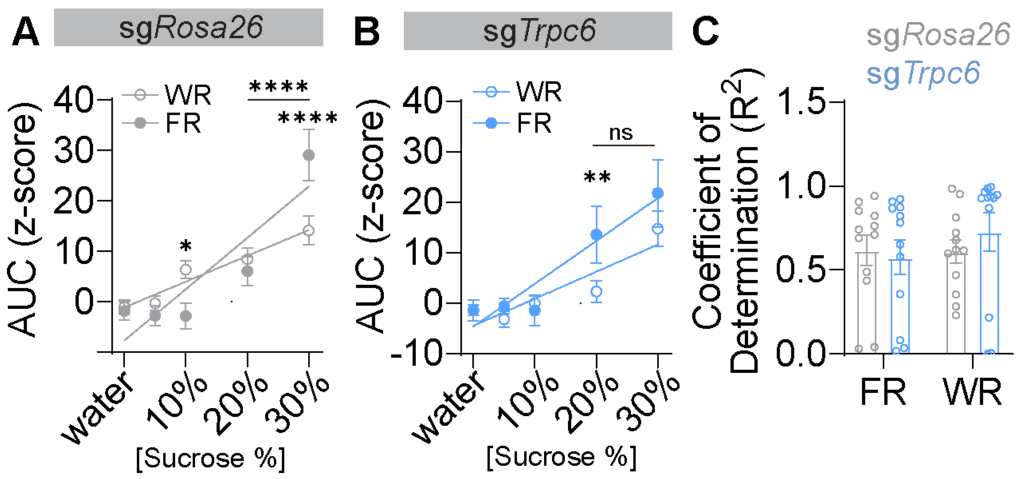

**Supplemental Figure 3.** Differential calcium responses following the solution access period in sg*Rosa26* and sg*Trpc6* mice. (A-B) GCaMP6m signal AUC of 10s post access period in response to water and increasing sucrose in sg*Rosa26* (A) and sg*Trpc6* (B) mice. In sg*Rosa26* mice (A), there are significantly larger responses to 10% sucrose under WR compared to FR and significantly larger responses to 30% sucrose in FR compared to WR. Like the AUC for the access period, the AUC for the post access period is larger at 30% compared to 20% under FR. In sg*Trpc6* mice (B) there are significantly larger responses to 20% sucrose FR compared to WR. However, unlike sg*Rosa26* mice, there is no significant difference in the AUC for the post access period at 30% compared to 20% under FR. (A-B) Two-way repeated measures ANOVA within subject comparison (*P<0.05, **P<0.01, and ****P<0.0001, Sidak post hoc comparison). (C) Linear regression analysis for individual mice reveals no significant differences between sg*Rosa26* and sg*Trpc6* mice under FR versus WR. Data are presented as mean ± S.E.M.

### Extended Data Table 1.

| Figure | | Test | N | Statistics | P | Posttest | p (post) |
|---|---|---|---|---|---|---|---|
| **Main Figure Statistics** | | | | | | | |
| 1 | A | Two-way RM ANOVA | N=7 mice | Interaction: F(6,72)=9.203; Effect of channel type: F(6,72)=15.20; Effect of group: F(1,12)=20.86 | P<0.0001, P<0.0001, P=0.0006 | Bonferroni multiple comparisons | ****p<0.0001 |
| | G | *AUC* D'Agostino & Pearson Test | $N_{sg\mathit{Rosa26}}$=4 mice, $n_{sg\mathit{Rosa26}}$=31 cells; $N_{sg\mathit{Trpc6}}$=3 mice, $n_{sg\mathit{Trpc6}}$=46 cells | $K^2_{sg\mathit{Rosa26}}=6.614$, $K^2_{sg\mathit{Trpc6}}=13.71$ | $P_{sg\mathit{Rosa26}}<0.05$, $P_{sg\mathit{Trpc6}}<0.01$ | | |
| | | *AUC* Mann—Whitney | $N_{sg\mathit{Rosa26}}$=4 mice, $n_{sg\mathit{Rosa26}}$=31 cells; $N_{sg\mathit{Trpc6}}$=3 mice, $n_{sg\mathit{Trpc6}}$=46 cells | U=412 | P<0.01 | | |
| | | *Peak* D'Agostino & Pearson Test | $N_{sg\mathit{Rosa26}}$=4 mice, $n_{sg\mathit{Rosa26}}$=31 cells; $N_{sg\mathit{Trpc6}}$=3 mice, $n_{sg\mathit{Trpc6}}$=46 cells | $K^2_{sg\mathit{Rosa26}}=11.78$, $K^2_{sg\mathit{Trpc6}}=33.68$ | $P_{sg\mathit{Rosa26}}<0.01$, $P_{sg\mathit{Trpc6}}<0.0001$ | | |
| | | *Peak* Mann—Whitney | $N_{sg\mathit{Rosa26}}$=4 mice, $n_{sg\mathit{Rosa26}}$=31 cells; $N_{sg\mathit{Trpc6}}$=3 mice, $n_{sg\mathit{Trpc6}}$=46 cells | U=378 | P<0.001 | | |
| | H | *Proportion* Chi-square | $\mathrm{Nonresponder}_{sg\mathit{Rosa26}}$=4 cells, $\mathrm{Responder}_{sg\mathit{Rosa26}}$=27 cells; $\mathrm{Nonresponder}_{sg\mathit{Trpc6}}$=21 cells, $\mathrm{Responder}_{sg\mathit{Trpc6}}$=25 cells | $\chi^2=9.059$, df=1, N=77 | P<0.01 | | |
| | J | *AUC* D'Agostino & Pearson Test | $N_{sg\mathit{Rosa26}}$=3 mice, $n_{sg\mathit{Rosa26}}$=36 cells; $N_{sg\mathit{Trpc6}}$=3 mice, $n_{sg\mathit{Trpc6}}$=47 cells | $K^2_{sg\mathit{Rosa26}}=13.52$, $K^2_{sg\mathit{Trpc6}}=53.47$ | $P_{sg\mathit{Rosa26}}<0.01$, $P_{sg\mathit{Trpc6}}<0.0001$ | | |
| | | *AUC* Mann—Whitney | $N_{sg\mathit{Rosa26}}$=3 mice, $n_{sg\mathit{Rosa26}}$=36 cells; $N_{sg\mathit{Trpc6}}$=3 mice, $n_{sg\mathit{Trpc6}}$=47 cells | U=598 | P<0.05 | | |

| | | | | | |
|---|---|---|---|---|---|
| | | *Peak* D'Agostino & Pearson Test | $N_{sg\mathit{Rosa26}}$=3 mice, $n_{sg\mathit{Rosa26}}$=36 cells; $N_{sg\mathit{Trpc6}}$=3 mice, $n_{sg\mathit{Trpc6}}$=47 cells | $K^2_{sg\mathit{Rosa26}}$=6.847, $K^2_{sg\mathit{Trpc6}}$=55.97 | $P_{sg\mathit{Rosa26}}<0.05$, $P_{sg\mathit{Trpc6}}<0.0001$ |
| | | *Peak* Mann—Whitney | $N_{sg\mathit{Rosa26}}$=3 mice, $n_{sg\mathit{Rosa26}}$=36 cells; $N_{sg\mathit{Trpc6}}$=3 mice, $n_{sg\mathit{Trpc6}}$=47 cells | U=523 | P<0.01 |
| | K | *Proportion* Chi-square | $Nonresponder_{sg\mathit{Rosa26}}$=10 cells, $Responder_{sg\mathit{Rosa26}}$=26 cells; $Nonresponder_{sg\mathit{Trpc6}}$=33 cells, $Responder_{sg\mathit{Trpc6}}$=14 cells | $\chi^2$=14.70, df=1, N=83 | P<0.0001 |
| 2 | A | *AUC* D'Agostino & Pearson Test | $N_{sg\mathit{Rosa26}}$=4 mice, $n_{sg\mathit{Rosa26}}$=34 cells; $N_{sg\mathit{Trpc6}}$=4 mice, $n_{sg\mathit{Trpc6}}$=33 cells | $K^2_{sg\mathit{Rosa26}}$=12.94, $K^2_{sg\mathit{Trpc6}}$=15.96 | $P_{sg\mathit{Rosa26}}<0.01$, $P_{sg\mathit{Trpc6}}<0.001$ |
| | | *AUC* Mann—Whitney | $N_{sg\mathit{Rosa26}}$=4 mice, $n_{sg\mathit{Rosa26}}$=34 cells; $N_{sg\mathit{Trpc6}}$=4 mice, $n_{sg\mathit{Trpc6}}$=33 cells | U=289 | P<0.001 |
| | | *Peak* D'Agostino & Pearson Test | $N_{sg\mathit{Rosa26}}$=4 mice, $n_{sg\mathit{Rosa26}}$=34 cells; $N_{sg\mathit{Trpc6}}$=4 mice, $n_{sg\mathit{Trpc6}}$=33 cells | $K^2_{sg\mathit{Rosa26}}$=2.610, $K^2_{sg\mathit{Trpc6}}$=3.770 | $P_{sg\mathit{Rosa26}}$=ns, $P_{sg\mathit{Trpc6}}$=ns |
| | | *Peak* Independent t-test | $N_{sg\mathit{Rosa26}}$=4 mice, $n_{sg\mathit{Rosa26}}$=34 cells; $N_{sg\mathit{Trpc6}}$=4 mice, $n_{sg\mathit{Trpc6}}$=33 cells | t=2.722, df=65 | P<0.01 |
| | | *Proportion* Chi-square | $Nonresponder_{sg\mathit{Rosa26}}$=6 cells, $Responder_{sg\mathit{Rosa26}}$=28 cells; $Nonresponder_{sg\mathit{Trpc6}}$=10 cells, $Responder_{sg\mathit{Trpc6}}$=23 cells | $\chi^2$=1.476, df=1, N=67 | P=ns |

| | | | | | |
|---|---|---|---|---|---|
| | B | Independent t-test | $N_{sg\mathit{Rosa26}}$=3 mice, $n_{sg\mathit{Rosa26}}$=290 cells; $N_{sg\mathit{Trpc6}}$=3 mice, $n_{sg\mathit{Trpc6}}$=512 cells | t=4.959, df=800 | P<0.0001 |
| | C | Independent t-test | $N_{sg\mathit{Rosa26}}$=3 mice, $n_{sg\mathit{Rosa26}}$=10 slices; $N_{sg\mathit{Trpc6}}$=3 mice, $n_{sg\mathit{Trpc6}}$=9 slices | t=3.153, df=17 | P<0.01 |
| | D | *Proportion* Fisher's exact test | Type $I_{sg\mathit{Rosa26}}$=66 cells, Type $II_{sg\mathit{Rosa26}}$=33 cells, Type $III_{sg\mathit{Rosa26}}$=1; Type $I_{sg\mathit{Trpc6}}$=89 cells, Type $II_{sg\mathit{Trpc6}}$=10 cells, Type $III_{sg\mathit{Trpc6}}$=1 cells | | P<0.001 |
| | E | *Frequency* D'Agostino & Pearson Test | $N_{sg\mathit{Rosa26}}$=3 mice, $n_{sg\mathit{Rosa26}}$=84 cells; $N_{sg\mathit{Trpc6}}$=3 mice, $n_{sg\mathit{Trpc6}}$=52 cells | $K^2_{sg\mathit{Rosa26}}$=14.42, $K^2_{sg\mathit{Trpc6}}$=10.16 | $P_{sg\mathit{Rosa26}}$<0.001, $P_{sg\mathit{Trpc6}}$<0.01 |
| | | *Frequency* Mann—Whitney | $N_{sg\mathit{Rosa26}}$=3 mice, $n_{sg\mathit{Rosa26}}$=84 cells; $N_{sg\mathit{Trpc6}}$=3 mice, $n_{sg\mathit{Trpc6}}$=52 cells | U=1452 | P<0.001 |
| | | *AUC* D'Agostino & Pearson Test | $N_{sg\mathit{Rosa26}}$=3 mice, $n_{sg\mathit{Rosa26}}$=84 cells; $N_{sg\mathit{Trpc6}}$=3 mice, $n_{sg\mathit{Trpc6}}$=52 cells | $K^2_{sg\mathit{Rosa26}}$=28.22, $K^2_{sg\mathit{Trpc6}}$=14.62 | $P_{sg\mathit{Rosa26}}$<0.0001, $P_{sg\mathit{Trpc6}}$<0.001 |
| | | *AUC* Mann—Whitney | $N_{sg\mathit{Rosa26}}$=3 mice, $n_{sg\mathit{Rosa26}}$=84 cells; $N_{sg\mathit{Trpc6}}$=3 mice, $n_{sg\mathit{Trpc6}}$=52 cells | U=2174 | P=ns |
| | F | *AUC* D'Agostino & Pearson Test | $N_{sg\mathit{Rosa26}}$=3 mice, $n_{sg\mathit{Rosa26}}$=40 cells; $N_{sg\mathit{Trpc6}}$=3 mice, $n_{sg\mathit{Trpc6}}$=30 cells | $K^2_{sg\mathit{Rosa26}}$=33.54, $K^2_{sg\mathit{Trpc6}}$=50.69 | $P_{sg\mathit{Rosa26}}$<0.0001, $P_{sg\mathit{Trpc6}}$<0.0001 |
| | | *AUC* Mann—Whitney | $N_{sg\mathit{Rosa26}}$=3 mice, $n_{sg\mathit{Rosa26}}$=40 cells; $N_{sg\mathit{Trpc6}}$=3 mice, $n_{sg\mathit{Trpc6}}$=30 cells | U=349 | P<0.01 |

| | | | | | |
|---|---|---|---|---|---|
| | | *Peak*<br>D'Agostino & Pearson Test | $N_{sgRosa26}$=3 mice, $n_{sgRosa26}$=40 cells; $N_{sgTrpc6}$=3 mice, $n_{sgTrpc6}$=30 cells | $K^2_{sgRosa26}$=6.054, $K^2_{sgTrpc6}$=39.26 | $P_{sgRosa26}$<0.05, $P_{sgTrpc6}$<0.0001 |
| | | *Peak*<br>Mann—Whitney | $N_{sgRosa26}$=3 mice, $n_{sgRosa26}$=40 cells; $N_{sgTrpc6}$=3 mice, $n_{sgTrpc6}$=30 cells | U=369 | P<0.01 |
| | | *Proportion*<br>Chi-square | $Nonresponder_{sgRosa26}$=11 cells, $Responder_{sgRosa26}$=29 cells; $Nonresponder_{sgTrpc6}$=15 cells, $Responder_{sgTrpc6}$=15 cells | $\chi^2$=3.717, df=1, N=70 | P<0.05 |
| | G | *AUC*<br>D'Agostino & Pearson Test | $N_{sgRosa26}$=3 mice, $n_{sgRosa26}$=27 cells; $N_{sgTrpc6}$=3 mice, $n_{sgTrpc6}$=32 cells | $K^2_{sgRosa26}$=29.76, $K^2_{sgTrpc6}$=20.39 | $P_{sgRosa26}$<0.0001, $P_{sgTrpc6}$<0.0001 |
| | | *AUC*<br>Mann—Whitney | $N_{sgRosa26}$=3 mice, $n_{sgRosa26}$=27 cells; $N_{sgTrpc6}$=3 mice, $n_{sgTrpc6}$=32 cells | U=393 | P=ns |
| | | *Peak*<br>D'Agostino & Pearson Test | $N_{sgRosa26}$=3 mice, $n_{sgRosa26}$=27 cells; $N_{sgTrpc6}$=3 mice, $n_{sgTrpc6}$=32 cells | $K^2_{sgRosa26}$=32.50, $K^2_{sgTrpc6}$=14.76 | $P_{sgRosa26}$<0.0001, $P_{sgTrpc6}$<0.001 |
| | | *Peak*<br>Mann—Whitney | $N_{sgRosa26}$=3 mice, $n_{sgRosa26}$=27 cells; $N_{sgTrpc6}$=3 mice, $n_{sgTrpc6}$=32 cells | U=398 | P=ns |
| | | *Proportion*<br>Chi-square | $Nonresponder_{sgRosa26}$=4 cells, $Responder_{sgRosa26}$=23 cells; $Nonresponder_{sgTrpc6}$=8 cells, $Responder_{sgTrpc6}$=24 cells | $\chi^2$=0.9376, df=1, N=59 | P=ns |

| | | | | | |
|---|---|---|---|---|---|
| | H | *AUC* D'Agostino & Pearson Test | $N_{sg\mathit{Rosa26}}$=3 mice, $n_{sg\mathit{Rosa26}}$=20 cells; $N_{sg\mathit{Trpc6}}$=4 mice, $n_{sg\mathit{Trpc6}}$=37 cells | $K^2_{sg\mathit{Rosa26}}$=13.89, $K^2_{sg\mathit{Trpc6}}$=46.10 | $P_{sg\mathit{Rosa26}}<0.01$, $P_{sg\mathit{Trpc6}}<0.0001$ |
| | | *AUC* Mann—Whitney | $N_{sg\mathit{Rosa26}}$=3 mice, $n_{sg\mathit{Rosa26}}$=20 cells; $N_{sg\mathit{Trpc6}}$=4 mice, $n_{sg\mathit{Trpc6}}$=37 cells | U=313 | P=ns |
| | | *Peak* D'Agostino & Pearson Test | $N_{sg\mathit{Rosa26}}$=3 mice, $n_{sg\mathit{Rosa26}}$=20 cells; $N_{sg\mathit{Trpc6}}$=4 mice, $n_{sg\mathit{Trpc6}}$=37 cells | $K^2_{sg\mathit{Rosa26}}$=6.594, $K^2_{sg\mathit{Trpc6}}$=30.73 | $P_{sg\mathit{Rosa26}}<0.05$, $P_{sg\mathit{Trpc6}}<0.0001$ |
| | | *Peak* Mann—Whitney | $N_{sg\mathit{Rosa26}}$=3 mice, $n_{sg\mathit{Rosa26}}$=20 cells; $N_{sg\mathit{Trpc6}}$=4 mice, $n_{sg\mathit{Trpc6}}$=37 cells | U=251.5 | P<0.05 |
| | | *Proportion* Chi-square | $Nonresponder_{sg\mathit{Rosa26}}$=3 cells, $Responder_{sg\mathit{Rosa26}}$=17 cells; $Nonresponder_{sg\mathit{Trpc6}}$=8 cells, $Responder_{sg\mathit{Trpc6}}$=29 cells | $\chi^2$=0.3655, df=1, N=57 | P=ns |
| 3 | B | *Proportion* Fisher's exact test | $Spontaneous_{sg\mathit{Rosa26}}$=42 cells, $Inactive_{sg\mathit{Rosa26}}$=34 cells; $Spontaneous_{sg\mathit{Trpc6}}$=39 cells, $Inactive_{sg\mathit{Trpc6}}$=27 cells | | P=ns |
| | C | D'Agostino & Pearson Test | $N_{sg\mathit{Rosa26}}$=5 mice, $n_{sg\mathit{Rosa26}}$=65 cells; $N_{sg\mathit{Trpc6}}$=6 mice, $n_{sg\mathit{Trpc6}}$=80 cells | $K^2_{sg\mathit{Rosa26}}$=42.53, $K^2_{sg\mathit{Trpc6}}$=39.38 | $P_{sg\mathit{Rosa26}}<0.0001$, $P_{sg\mathit{Trpc6}}<0.0001$ |
| | | Mann—Whitney | $N_{sg\mathit{Rosa26}}$=5 mice, $n_{sg\mathit{Rosa26}}$=65 cells; $N_{sg\mathit{Trpc6}}$=6 mice, $n_{sg\mathit{Trpc6}}$=80 cells | U=2445 | P=ns |

| | | | | | | | |
|---|---|---|---|---|---|---|---|
| | D | D'Agostino & Pearson Test | $N_{sgRosa26}$=5 mice, $n_{sgRosa26}$=51 cells; $N_{sgTrpc6}$=6 mice, $n_{sgTrpc6}$=56 cells | $K^2_{sgRosa26}$=46.31 , $K^2_{sgTrpc6}$=85.21 | $P_{sgRosa26}$<0.0001, $P_{sgTrpc6}$<0.0001 | | |
| | | Mann—Whitney | $N_{sgRosa26}$=5 mice, $n_{sgRosa26}$=51 cells; $N_{sgTrpc6}$=6 mice, $n_{sgTrpc6}$=56 cells | U=1377 | P=ns | | |
| | E | D'Agostino & Pearson Test | $N_{sgRosa26}$=5 mice, $n_{sgRosa26}$=65 cells; $N_{sgTrpc6}$=6 mice, $n_{sgTrpc6}$=67 cells | $K^2_{sgRosa26}$=7.325 , $K^2_{sgTrpc6}$=6.143 | $P_{sgRosa26}$<0.05, $P_{sgTrpc6}$<0.05 | | |
| | | Mann—Whitney | $N_{sgRosa26}$=5 mice, $n_{sgRosa26}$=65 cells; $N_{sgTrpc6}$=6 mice, $n_{sgTrpc6}$=67 cells | U=2134 | P=ns | | |
| | F | D'Agostino & Pearson Test | $N_{sgRosa26}$=8 mice, $n_{sgRosa26}$=28 cells; $N_{sgTrpc6}$=3 mice, $n_{sgTrpc6}$=36 cells | $K^2_{sgRosa26}$=10.16 , $K^2_{sgTrpc6}$=1.330 | $P_{sgRosa26}$<0.01, $P_{sgTrpc6}$=ns | | |
| | | Mann—Whitney | $N_{sgRosa26}$=8 mice, $n_{sgRosa26}$=28 cells; $N_{sgTrpc6}$=3 mice, $n_{sgTrpc6}$=36 cells | U=473 | P=ns | | |
| | H | Two-way RM ANOVA | $N_{sgRosa26}$=13 mice, $n_{sgRosa26}$=15 cells; $N_{sgTrpc6}$=9 mice, $n_{sgTrpc6}$=26 cells | Effect of step: F(1.893, 73.83); Effect of subject: F(39, 897) = 49.89 | Effect of step: P<0.0001; Effect of subject: P<0.0001 | Bonferroni's multiple comparisons | p=ns |
| 4 | D | Two-way RM ANOVA | $N_{sgRosa26}$=12 mice, $N_{sgTrpc6}$=12 mice | Effect of spout ID: F(4,88) = 43.02; Effect of subject: F(22,88) = 7.247 | Effect of spout ID: P<0.0001; Effect of subject: P<0.0001 | Tukey's multiple comparisons | *p<0.05, **p<0.01, ***p<0.001, ****p<0.0001 |
| | E | Independent t-test | $N_{sgRosa26}$=12 mice, $N_{sgTrpc6}$=12 mice | t=0.9955, df=22 | P=ns | | |

| | | | | | | | |
|---|---|---|---|---|---|---|---|
| | G | Two-way RM ANOVA | $N_{sg\textit{Rosa26}}$=12 mice, $N_{sg\textit{Trpc6}}$=12 mice | Interaction: F(4,88) = 3.413; Effect of spout ID: F(4,88) = 174.2; Effect of subject: F(22,88) = 7.311 | Interaction: P=0.0121 Effect of spout ID: P<0.0001; Effect of subject: P<0.0001 | Tukey's multiple comparisons | *p<0.05, **p<0.01, ***p<0.001, ****p<0.0001 |
| | H | Independe nt t-test | $N_{sg\textit{Rosa26}}$=12 mice, $N_{sg\textit{Trpc6}}$=12 mice | t=2.823, df=22 | P<0.01 | | |
| 5 | B | Two-way RM ANOVA | $N_{sg\textit{Rosa26}}$=12 mice, $N_{sg\textit{Trpc6}}$=12 mice | Effect of spout ID: F(4,88) = 41.87; Effect of subject: F(22,88) = 4.052 | Effect of spout ID: P<0.0001; Effect of subject: P<0.0001 | Tukey's multiple comparisons | *p<0.05, **p<0.01, ***p<0.001, ****p<0.0001 |
| | C | Independe nt t-test | $N_{sg\textit{Rosa26}}$=12 mice, $N_{sg\textit{Trpc6}}$=12 mice | t=0.8182, df=22 | P=ns | | |
| | E | Two-way RM ANOVA | $N_{sg\textit{Rosa26}}$=12 mice, $N_{sg\textit{Trpc6}}$=12 mice | Effect of spout ID: F(4,88) = 49.11; Effect of subject: F(22,88) = 3.362 | Effect of spout ID: P<0.0001; Effect of subject: P<0.0001 | Tukey's multiple comparisons | *p=0.0505, **p<0.01, ***p<0.001, ****p<0.0001 |
| | F | Independe nt t-test | $N_{sg\textit{Rosa26}}$=12 mice, $N_{sg\textit{Trpc6}}$=12 mice | t=1.284, df=22 | P=ns | | |
| | G | *Water* Paired t-test | $N_{WR}$=12 mice; $N_{FR}$=12 mice | t = 3.647, df = 11 | P<0.01 | | |

| | | | | | |
|---|---|---|---|---|---|
| | | *5% sucrose* Paired t-test | $N_{WR}$=12 mice; $N_{FR}$=12 mice | t = 5.608, df = 11 | P<0.001 |
| | | *10% sucrose* Paired t-test | $N_{WR}$=12 mice; $N_{FR}$=12 mice | t = 4.546, df = 11 | P<0.001 |
| | | *20% sucrose* Paired t-test | $N_{WR}$=12 mice; $N_{FR}$=12 mice | t = 0.6625, df = 11 | P=ns |
| | | *30% sucrose* Paired t-test | $N_{WR}$=12 mice; $N_{FR}$=12 mice | t = 2.439, df = 11 | P<0.05 |
| | H | *Water* Paired t-test | $N_{WR}$=12 mice; $N_{FR}$=12 mice | t = 0.5427, df = 11 | P=ns |
| | | *5% sucrose* Paired t-test | $N_{WR}$=12 mice; $N_{FR}$=12 mice | t = 2.206, df = 11 | P<0.05 |
| | | *10% sucrose* Paired t-test | $N_{WR}$=12 mice; $N_{FR}$=12 mice | t = 1.955, df = 11 | P=ns |
| | | *20% sucrose* Paired t-test | $N_{WR}$=12 mice; $N_{FR}$=12 mice | t = 1.681, df = 11 | P=ns |
| | | *30% sucrose* Paired t-test | $N_{WR}$=12 mice; $N_{FR}$=12 mice | t = 0.01657, df = 11 | P=ns |
| **Supplementary Data Statistics** | | | | | |
| 1 | D | Independent t-test | $N_{sgRosa26}$=12 mice, $N_{sgTrpc6}$=12 mice | t=1.113, df=22 | P=ns |

| | | | | | | | |
|---|---|---|---|---|---|---|---|
| 2 | A | sg*Rosa26* Two-way RM ANOVA, within group comparisons | $N_{WR}$=12 mice; $N_{FR}$=12 mice | Interaction: F(4, 44)=6.388; Effect of spout ID: F(4, 44) = 96.08; Effect of restriction: F(1, 11)=24.25 | Interaction: P=0.0004; Effect of spout ID: P<0.0001; Effect of restriction: P=0.0005 | Sidak's multiple comparisons | *p<0.05, **p<0.01, ***p<0.001, ****p<0.0001 |
| | B | sg*Trpc6* Two-way RM ANOVA, within group comparisons | $N_{WR}$=12 mice; $N_{FR}$=12 mice | Interaction: F(4, 44)=19.65; Effect of spout ID: F(4, 44) = 75.86 | Interaction: P<0.0001; Effect of spout ID: P<0.0001 | Sidak's multiple comparisons | *p<0.05, **p<0.01, ***p<0.001, ****p<0.0001 |
| | C | *Regression Coefficient* Two-way RM ANOVA | $N_{sgRosa26}$=12 mice, $N_{sgTrpc6}$=12 mice | Effect of restriction: F(1,22) = 21.36 | P=0.0001 | Uncorrected Fisher's LSD | **p<0.01 |
| 3 | A | sg*Rosa26* Two-way RM ANOVA, within group comparisons | $N_{WR}$=12 mice; $N_{FR}$=12 mice | Interaction: F(4, 44)=9.717; Effect of spout ID: F(4, 44) = 23.36 | Interaction: P<0.0001; Effect of spout ID: P<0.0001 | Sidak's multiple comparisons | *p<0.05, **p<0.01, ***p<0.001, ****p<0.0001 |
| | B | sg*Trpc6* Two-way RM ANOVA, within group comparisons | $N_{WR}$=12 mice; $N_{FR}$=12 mice | Interaction: F(4, 44)=3.110; Effect of spout ID: F(4, 44) = 12.77 | Interaction: P=0.0244; Effect of spout ID: P<0.0001 | Sidak's multiple comparisons | *p<0.05, **p<0.01, ***p<0.001, ****p<0.0001 |
| | C | *Regression Coefficient* Two-way RM ANOVA | $N_{sgRosa26}$=12 mice, $N_{sgTrpc6}$=12 mice | | P=ns | Sidak's multiple comparisons | P=ns |
